# Ongoing coevolution between reintroduced *Phengaris teleius* butterflies and their *Myrmica* host ants

**DOI:** 10.64898/2025.12.02.691951

**Authors:** Daniel Sánchez-García, Irma Wynhoff, Patrizia d’Ettorre, Chloé Leroy, Joanna Kajzer-Bonk, István Elek Maák, Francesca Barbero, Luca Pietro Casacci, Magdalena Witek

**Author notes:** Correspondence: Daniel Sánchez-García < / >. Raw data and analysis scripts (R code) supporting this manuscript are publicly available via Zenodo (https://doi.org/10.5281/zenodo.17791047, version 1.1).

## Abstract

Coevolutionary interactions between parasites and hosts are key drivers of biological adaptation. In this study, we explore the evolutionary response of the social parasitic butterfly *Phengaris teleius* to its host ant, *Myrmica scabrinodis*, taking advantage of a unique opportunity: the reintroduction of the butterfly in the Netherlands thirty years ago. We compared the degree of host mimicry and behavioural performance of caterpillars between the reintroduced and the Polish source population. After about thirty generations, chemical and vibroacoustical signal profiles have diverged. Chemical mimicry remained limited during the pre-adoption phase for both groups; however, in the post-adoption phase, the source population showed significantly higher chemical similarity to their local hosts. In contrast, reintroduced pre-adoption caterpillars evolved vibroacoustic signals closely resembling local hosts, also resulting in a stronger response from their local host ants. This suggests that adoption is driven by acoustics and subsequently serves as a selective filter promoting post-entry chemical refinement. Behavioural data evince that despite the differences between the different communication channels, the combination of signals remains sufficient to ensure recognition and integration in both host-parasite systems. These results illustrate how social parasites involved in multisensory mimicry can rapidly recalibrate strategies to remain functional in new ecological contexts.

## Introduction

Interactions between species, including coevolutionary dynamics between parasites and hosts, represent critical forces that drive evolution and speciation [1,2]. A peculiar type of antagonistic interaction, where the parasite exploits a whole society instead of a single organism, is called social parasitism. It occurs in diverse taxa, including birds, fish, and particularly, social insects like bees or ants [3–5]. Several strategies evolved by social parasites of ants allow them to infiltrate and integrate within host colonies by disrupting ant communication system [6]. Among the most intriguing adaptations are those employed by caterpillars of *Phengaris* butterflies to exploit resources within *Myrmica* ant colonies. The caterpillars of these obligate social parasites are retrieved by *Myrmica* foraging ants and carried inside the colony, where they live until adult emergence [7]. *Phengaris* immature instars “cheat” their hosts, making ants firstly adopt and then tolerate them, by corrupting ants’ primary communication channels based on chemical [8–10] and vibroacoustic signals [11,12].

Nestmate recognition in social insects is mainly mediated by cuticular hydrocarbons (CHCs) [13]. *Myrmica* ants possess species-specific CHC profiles that can vary both qualitatively and quantitatively, with intraspecific variation observed among populations from distinct geographical localities [14–16]. Sharing a similar chemical profile enables ants of the same colony to discriminate between nestmates and intruders [17]. However, the early stage of *Phengaris* caterpillars (pre-adoption) possesses a simple blend of CHCs that mimics those of *Myrmica* ant brood, thus deceiving the foragers and promoting the adoption of the parasites within host colonies [8]. Soon after entering the nest (post-adoption), the chemical profiles of the caterpillars change and resemble more those of the ant colony members, allowing the parasites to live in the host society until pupation [9]. Besides imitating chemical recognition cues, *Phengaris* caterpillars can also produce “calls” that closely resemble the vibroacoustic signals emitted by *Myrmica* ants, especially queens [18–20]. Since vibrations are used for inter- and intra-cast communication in ants, the queen-like signals produced by the parasite effectively deceive the workers that treat the caterpillars as valuable items in the colony hierarchy [18].

Different *Phengaris* species and populations use various *Myrmica* ants as hosts, and local adaptations between these butterflies and their host ants have been observed in some populations of the ‘cuckoo’-feeding *Phengaris* group [10,16,21,22]. Cuckoo species are fed directly by nurse ants via trophallaxis; thus, they require spotless integration and full acceptance as nestmate in the host society, possibly leading to high host-specific populations [23]. In contrast, predatory species like *Phengaris arion* [24] or *Phengaris teleius* [25], actively prey on ant broods and, once they are retrieved from outside into the nest, they only need not to be discarded as intruders, having the possibility to parasitize a wider range of hosts.

To test whether generalist social parasites like *P. teleius* could evolve adaptations to their local hosts and to detect potential ongoing coevolutionary processes in this host-parasite system, we surveyed a reintroduced and its paired source population. Briefly, 86 *P. teleius* adult butterflies were moved from the Polish population occurring in Krakow to the Dutch reserve of Moerputten in 1990 [more details in 26] and the successful reintroduction led to the establishment of a population in the Netherlands [27]. A gap of almost 30 butterfly generations since the reintroduction offered a unique opportunity to study the chemical and acoustical adaptations of *P. teleius* butterflies to their main and most abundant host ant, *Myrmica scabrinodis* [28]. We hypothesized that: 1) the CHC profiles and 2) the vibroacoustic signals of the butterfly caterpillars differ between the two populations and are more similar to the signals of the local *M. scabrinodis* host ants; 3) the vibroacoustic signals produced by the social parasites are able to stimulate more benevolent behaviours in their sympatric host ants; and 4) the caterpillars exposed to host ants from sympatric populations undergo a more successful adoption process and show a higher survival within the local ant colonies.

## Material and Methods

### Collection of Phengaris teleius caterpillars and Myrmica scabrinodis ants

*P. teleius* caterpillars and *M. scabrinodis* ant colonies were collected from the source population (PL) in the Vistula River Valley in Krakow, Poland (50°01′N, 19°54′E) and from the reintroduced population (NL) in the nature reserve of Moerputten in the Netherlands (51°41′N, 5°15′E). The samples to study the pre-adoption CHC profile and vibroacoustic signals were collected in August 2019; the samples to perform the behavioural assay and study the post-adoption CHC profile were collected in July and August 2020; and the samples to perform the playback experiment were collected in August and October 2021.

*P. teleius* caterpillars were collected from the flowers of their foodplant, *Sanguisorba officinalis* [29]. These samples are hereafter labeled as “pre-adoption”. *M. scabrinodis* ant workers were collected and housed in plastic containers until used for different experiments. Before adopting a *P. teleius* caterpillar, we called them laboratory nests; when they hosted a caterpillar, they were named “post-adoption” colonies. In the same way, once inside the nests, *P. teleius* caterpillars were referred to as “post-adoption” caterpillars. For detailed collection and transport details see *Methods S1*.

### CHC extraction and GC-MS analysis

CHC compounds of pre-adoption *P. teleius* caterpillars were extracted within a few hours after they left their host plant. Pre-adoption *M. scabrinodis* workers were processed one day after collecting ants in the field. Ant samples consisted of a group of five ants pooled together into the same vial. Three replicates were taken from each sampled colony. Pre-adoption samples were extracted from a total of 21 and 24 *M. scabrinodis* and 25 and 19 *P. teleius* caterpillars from Poland and the Netherlands, respectively. The samples of post-adopted *P. teleius* caterpillars and their host *M. scabrinodis* were collected during the behavioural experiment on caterpillar adoption, whose protocol is described below. The CHC profiles of the caterpillars and ants were extracted three days after caterpillar adoption. For CHC extraction, we collected a total of three *P. teleius* caterpillars from the Polish source population adopted by their sympatric host ants and three adopted by the Dutch host ants; two *P. teleius* caterpillars from the Dutch population adopted by their sympatric host ants and two adopted by Polish ants. We also extracted the CHC profile from these host ants.

CHCs were extracted from individual caterpillars and pools of five ant workers (killed by freezing) by placing them in a glass vial with 200 μL of hexane for 10 minutes. After extraction, all vials (from pre- and post-adopted experiments) were stored at −20 °C until analysis. Prior analysis an internal standard was added to the extract. A detailed protocol with a description of subsequent parts of sample processing is presented in *Methods S1*.

### CHC raw data processing

The chromatograms were analyzed in MSD ChemStation E.02.01.1177 (Agilent Technologies) with the RTE integrator to calculate the area of each peak of interest using the proportion of the sum over the area of all peaks applying the parameter values presented in *Methods S1*. Hydrocarbons were identified based on their mass spectra and retention times, and compared with known standards. CHC peaks retention time was used to perform an automatic aligning of the compounds by applying an alignment algorithm implemented in the *align_chromatograms()* function [30]. After aligning, all peaks were double-checked to ensure a correct identification. Only peaks with a retention time from 8 to 26.5 minutes were selected. Alkanes were removed from the dataset because of sample contamination to reduce noise during the analysis. Therefore, our CHC dataset was composed of only methylated alkanes and alkenes, which still perfectly define our goal, since these compounds are the primary drivers of nestmate recognition [13].

CHC intensity values after peak integration were transformed to absolute abundance data by correcting with the internal standard. Data were filtered by selecting the compounds with an absolute abundance higher than or equal to 0.02 % per sample. Only the compounds occurring in at least 40 % of the samples per group (species+population) were taken. CHC absolute abundance was then divided by the dry mass of the sample, and log_10_(*x*) was applied to avoid over-representation of very abundant compounds. Ant and caterpillar bodies were dried at 50 °C for a week, and subsequently weighed in a Radwag microbalance MYA 5.4Y (±1 μg) to estimate their dry mass.

### CHC profile statistical analysis

CHC profile distances between groups of individuals characterized by species (*P. teleius* caterpillar and *M. scabrinodis* ant), adoption state (pre- and post-adoption) and population (Poland and the Netherlands) were calculated using *vegdist()* [31] with the Bray-Curtis dissimilarity index. Differences among these groups were assessed via permutational analysis of variance (PERMANOVA) using *adonis2()* [31]. The Bray-Curtis distance matrix was subsetted in three different ways: 1) pre-adoption *P. teleius* and pre-adoption *M. scabrinodis*, 2) post-adoption *P teleius* and post-adoption *M. scabrinodis* and 3) pre-adoption *P. teleius* and post-adoption *P. teleius*. From each subset, distances were extracted and fitted to a generalised linear mixed model (GLMM) with Gaussian distribution. We used caterpillar and ant populations as fixed effects, while the pairs of samples used to compute distances were included as random effects to account for non-independence using *glmmTMB()* [32,33]. Predictor significance was tested with ANOVA using *Anova()* [34] and the pairwise comparisons were performed via estimated marginal means (EMMs) with Holm correction using *emmeans()* [35]. To assess differences between groups at the CHC compound level we employed a series of generalised linear models (GLMs) with a binomial distribution. To mitigate potential bias from separation (perfect prediction due to large compound abundance differences between groups) and ensure robust coefficient estimates in our binomial GLMs, we used the brglmFit method in *glm()* [36,37], which implements bias reduction techniques. For each compound, we fitted a GLM that was constructed to evaluate the probability of belonging to the comparison level based on the compound abundance relative to the reference level (intercept). The magnitude of the coefficient (*β*_1_) associated with the comparison level reflects the strength of the effect. This coefficient quantifies the change in the log-odds of compound abundance when comparing the comparison level to the reference level. A positive coefficient indicates an increase in compound abundance relative to the reference, while a negative coefficient indicates a decrease. We compared the coefficients obtained from each individual GLM. This allowed us to order the compounds based on the magnitude and direction of their response.

### Vibroacoustics signal recordings

We utilized custom-made equipment designed for recording the sounds produced by undisturbed and unstressed insects [38]. We recorded a total of eight caterpillars, five ant queens and 10 ant workers from Poland, and six caterpillars, three ant queens and 10 ant workers from the Netherlands. *P. teleius* caterpillars and ants were individually positioned on the microphone surface inside the recording chamber and recorded in the morning under room temperature conditions (23–25 °C). Recording sessions lasted 10 minutes, starting 5 minutes after introducing the specimens into the recording chamber. We processed the vibroacoustic signals and measured six parameters. For detailed protocols regarding signal processing, software specifications, and definitions of the vibroacoustic variables measured, refer to the *Methods S1*.

### Playback experiments

To test whether the sounds emitted by the caterpillars were able to produce a greater behavioural response in their sympatric host ants, we conducted playback experiments using four colonies from Poland and five colonies from the Netherlands in August and October 2021. Playback experiments were carried out in artificial arenas made of plastic cylinders (7 × 7 × 5 cm). A speaker was glued to the bottom of each arena. We covered the speaker with a thin layer of slightly damp soil to simulate natural conditions. In each arena, we introduced five *M. scabrinodis* ant workers and allowed them to settle for 10 minutes before exposing them to one of the five vibroacoustic signals previously recorded and emitted by 1) *M. scabrinodis* queens, 2) *M. scabrinodis* workers, 3) sympatric caterpillars of *P. teleius* and 4) allopatric caterpillars of *P. teleius*; a 5) white noise was used as a control. We employed MP3 devices to play continuous loops of the original recordings, adjusting the volume to match the natural level (for detailed volume adjustments, refer to [12]). Each experimental trial spanned 30 minutes, with behavioural observations conducted in one-minute intervals for each arena, in sequential order for the six signal types, totaling six minutes of signal exposure per trial. During trials, we observed five different behaviours, i.e. walking, staying, antennating, guarding and digging, previously described [12,38]. We replicated the playback experiment per each colony from two to three times, using new *Myrmica* ants for each arena. The signal source in each arena was randomized to account for potential positional effects. Prior to each trial, new soil was introduced, and all equipment was thoroughly cleaned with absolute ethanol.

### Vibroacoustics data analysis

We used the normalized vibroacoustic parameters derived from the signal units emitted by the recorded specimens to compute a Euclidean distance matrix. This matrix was then used to perform a non-metric multidimensional scaling (NMDS) ordination using *metaMDS()* [31]. In order to assess disparities in vibroacoustic signals between the host (queens and workers) and pre-adoption parasitic caterpillars from the two populations, we carried out analysis of similarities (ANOSIM), conducting 9999 permutations. This analysis was conducted using *anosim()* [31]. The Euclidean distances were also analyzed separately for queens and workers. The data were fitted to GLMMs using *glmmTMB()* [32,33] with ant and caterpillar populations as fixed factors and the pair of samples used to compute the distances as random factors to account for non-independence. Differences in the average values of vibroacoustic parameters calculated on the signal units emitted by ant caste individuals and pre-adoption caterpillars from the two populations were tested with LMMs, incorporating individuals as a random factor by using *lmer()* [39]. Predictor significance was evaluated with ANOVA using *Anova()* [34] and post-hoc pairwise comparisons were performed using estimated marginal means (EMMs) tests with Holm correction via *emmeans()* [35].

We also compared the impact of the vibroacoustic signals on the total instances of behaviours exhibited by the worker ants of the two populations. We employed a negative binomial generalised linear mixed-effects model with “colony” as a random factor using *glmer.nb()* [39]. Predictor significance was tested with ANOVA using *Anova()* [34] and post-hoc pairwise comparisons were conducted using estimated marginal means (EMMs) tests with Bonferroni correction via *emmeans()* [35].

### Behavioural assay: adoption and survival of Phengaris teleius caterpillars

Behavioural experiments were performed in Warsaw in the laboratory of the Museum and Institute of Zoology in August 2020. For the experiments, we used 15 colonies from the Polish population and 11 colonies from the Dutch population. One sub-colony from each colony was used to observe the adoption of a caterpillar from the sympatric population, and another sub-colony to observe the adoption of an allopatric caterpillar. For detailed rearing and colony maintenance protocols, see *Methods S1*. Altogether, 15 Polish caterpillars were introduced to Polish ants and 11 to Dutch ants, while 15 Dutch caterpillars were introduced to Polish ants and 11 to Dutch ants. Only fourth-instar caterpillars were used. To observe adoption, a caterpillar was introduced into the foraging area, and detailed observations were made for 60 minutes following the first contact with workers. We recorded all behavioural interactions (categorized as inspection, positive, or negative behaviours) and the number of sugary drops secreted by the caterpillars. If adoption did not happen during the first 60 minutes, we monitored the colonies to determine if adoption occurred within 24 hours, and checked caterpillar survival daily. For detailed observation protocols and behavioural definitions, see *Methods S1*.

### Behavioural data analysis

Behavioural observations could be underestimated when caterpillars were adopted or killed before the end of the observation period; therefore, antennation events, positive behaviours, negative behaviours, and drops were standardized by dividing counts by observation time (minutes) and multiplying by 60. Antennation event counts were fitted to a GLMM with Poisson distribution, with ant, caterpillar population and their interaction as predictors, and ant colony as random factor using *glmmTMB()* [32,33]. Positive and negative behaviours were fitted similarly using a negative binomial Type I and II distribution, respectively. The counts of drops produced by the caterpillars were also fitted to a GLMM with negative binomial Type II distribution. Predictor significance was tested with ANOVA using *Anova()* [34], and the groups were compared pairwise using estimated marginal means (EMMs) tests with Bonferroni correction via *emmeans()* [35]. The number of positive and negative behaviours was correlated with the number of drops produced by the caterpillars using *cor.test()* [37]. Adoption success was fitted to a GLMM with binomial distribution, with ant and caterpillar populations as predictors, and ant colony as random factor using *glmer()* [39]. Predictor significance was tested with ANOVA, followed by pairwise comparisons using EMMs with Bonferroni correction. Survival duration was analyzed with a Cox proportional-hazards model using *coxph()* [40] with ant and caterpillar population as predictors. Predictor significance was assesesd with ANOVA via *Anova()* [34]. Hazards ratios indicate a covariate level that is neutral (= 1), positively (< 1) or negatively (> 1) associated with the length of survival respecting the reference level. Predicted survival proportions were estimated with *survfit()* [40], and survival curves were pairwise compared via EMMs with Bonferroni correction. All statistical analyses were performed in R [37].

## Results

### Cuticular hydrocarbon adaptations

A total of 31 cuticular hydrocarbon compounds were identified both for *M. scabrinodis* ants and *P. teleius* caterpillars, and their presence and abundance in the different groups are presented in Figure S1. Additionally, a graphical representation of the CHC profiles can be found in Figure S2.

The cuticular hydrocarbon (CHC) profiles differed among all *M. scabrinodis* host ants and *P. teleius* caterpillars, both before and after adoption (d.f. = 3, F = 260.69, p = 0.001). Additionally, the population of origin of the individuals significantly influenced their CHC composition (d.f. = 1, F = 7.36, p = 0.002). However, not all groups displayed differences between their populations (d.f. = 3, F = 5.72, p = 0.001; Figure 1a).

**Figure 1.**
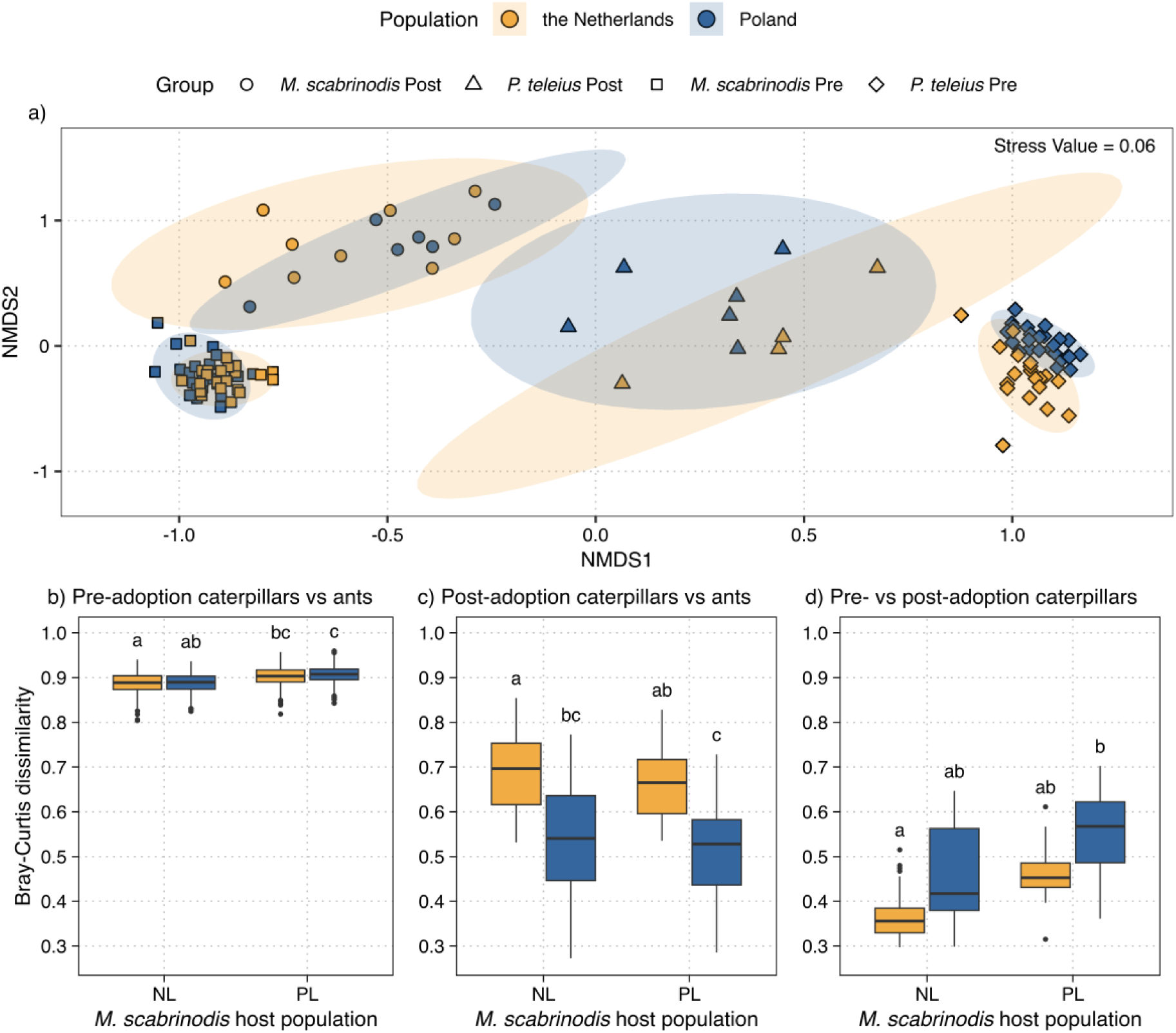
Comparison of CHC profiles between the reintroduced and source population of *P. teleius* caterpillars and their *M. scabrinodis* host ant. (a) NMDS based on Bray-Curtis distances (Stress = 0.04). Colors denote population: yellow (the Netherlands) and blue (Poland). Shapes denote species and stage: circle (post-adoption *M. scabrinodis*), triangle (post-adoption *P. teleius*), rectangle (pre-adoption *M. scabrinodis*) and cross (pre-adoption *P. teleius*). (b–d) Bray-Curtis dissimilarities between: (b) pre-adoption pairs, (c) post-adoption pairs, and (d) pre-vs. post-adoption caterpillars. The x-axis denotes the population of the host. The boxplot colors denote the population of origin for *P. teleius*: yellow (the Netherlands) and blue (Poland). Horizontal bars represent the median, boxes represent the 25% and 75% quartiles and whiskers the maximum and minimum values. Dots represent outliers. Different letters above boxplots denote pairwise significant differences based on estimated marginal means tests.

The differences in CHC profiles between pre-adoption *P. teleius* caterpillar and *M. scabrinodis* host ants were partially explained by the ant population (Table S1a). Dutch pre-adoption caterpillars had a CHC profile more similar to their local host ants (NL) than to the original hosts (PL). However, caterpillars from both populations showed a more similar profile to the Dutch host ants, with no significant differences between them (Figure 1b; Table S1b). Both caterpillar populations followed the same pattern regarding the abundance of CHCs with respect to the Dutch host ants with some compounds of lower abundance and some with higher one (Figure S3).

After adoption, the differences between the caterpillars and host ants were explained by the population of the caterpillars, but no effect of host population was detected like in the pre-adoption stage (Table S2a). Caterpillars from Poland showed a significantly more similar profile to both populations of the host ants in comparison to the Dutch caterpillars with the closest similarity to the ant CHC profile overall when parasitizing their sympatric hosts (Figure 1c; Table S2b). The compounds with lower absolute abundance in pre-adoption caterpillars than in the host ant showed no significant change after adoption. Similarly, most of the more abundant compounds did not show differences or the differences got reduced after caterpillar adoption, indicating that the caterpillar chemical profiles closely resembled the one of the host ant (Figure S3. and 4).

We also analyzed how much the CHC profile of the caterpillars changed during the adoption by host ants from different populations. The chemical signature of the caterpillars was significantly affected by the population of both post-adoption caterpillars and host ants (Table S3a). The CHC profile of *P. teleius* caterpillars from the Netherlands changed less than that of the Polish caterpillars. The Dutch caterpillars parasitizing their sympatric host ants showed the smallest changes in their CHC profile. In contrast, the Polish caterpillars reared by their sympatric host ants underwent the most significant changes in their CHC profile (Figure 1d; Table S3b). The differences that were observed between pre-adoption caterpillars were mostly quantitative and disappeared after adoption (Figure S5).

### Vibroacoustic adaptations

We recorded and analyzed the stridulations made by both worker and queen ants of *M. scabrinodis*, as well as the vibroacoustic signals produced by pre-adoption caterpillars of *P. teleius* collected in Poland and the Netherlands. Ant and caterpillar vibroacoustic signals consisted of sequences (trains) of variable numbers of units (as shown in Figure S6). Non-metric multidimensional scaling of the vibroacoustic patterns of ants and caterpillars showed a separation among populations and species (Figure 2) confirmed by an analysis of similarities (ANOSIM: R = 0.537, p < 0.001). The signals produced by *M. scabrinodis* queens were statistically distinct between populations (ANOSIM: R = 0.407, p < 0.001) as well as those emitted by workers (ANOSIM: R = 0.568, p < 0.001). Also, the vibroacoustic pattern of *P. teleius* pre-adoption caterpillars differed between populations (ANOSIM: R = 0.536, p < 0.001). The emissions of the parasitic caterpillars appeared significantly distinct from the host caste emissions, but *P. teleius* caterpillar signals were closer to those of queens. Nevertheless, in the source population, the signals of queens and *P. teleius* caterpillars showed the highest degree of similarity with an R value close to 0 (Poland, ANOSIM: R = 0.082, p = 0.016; the Netherlands, ANOSIM: R = 0.231, p < 0.001).

**Figure 2.**
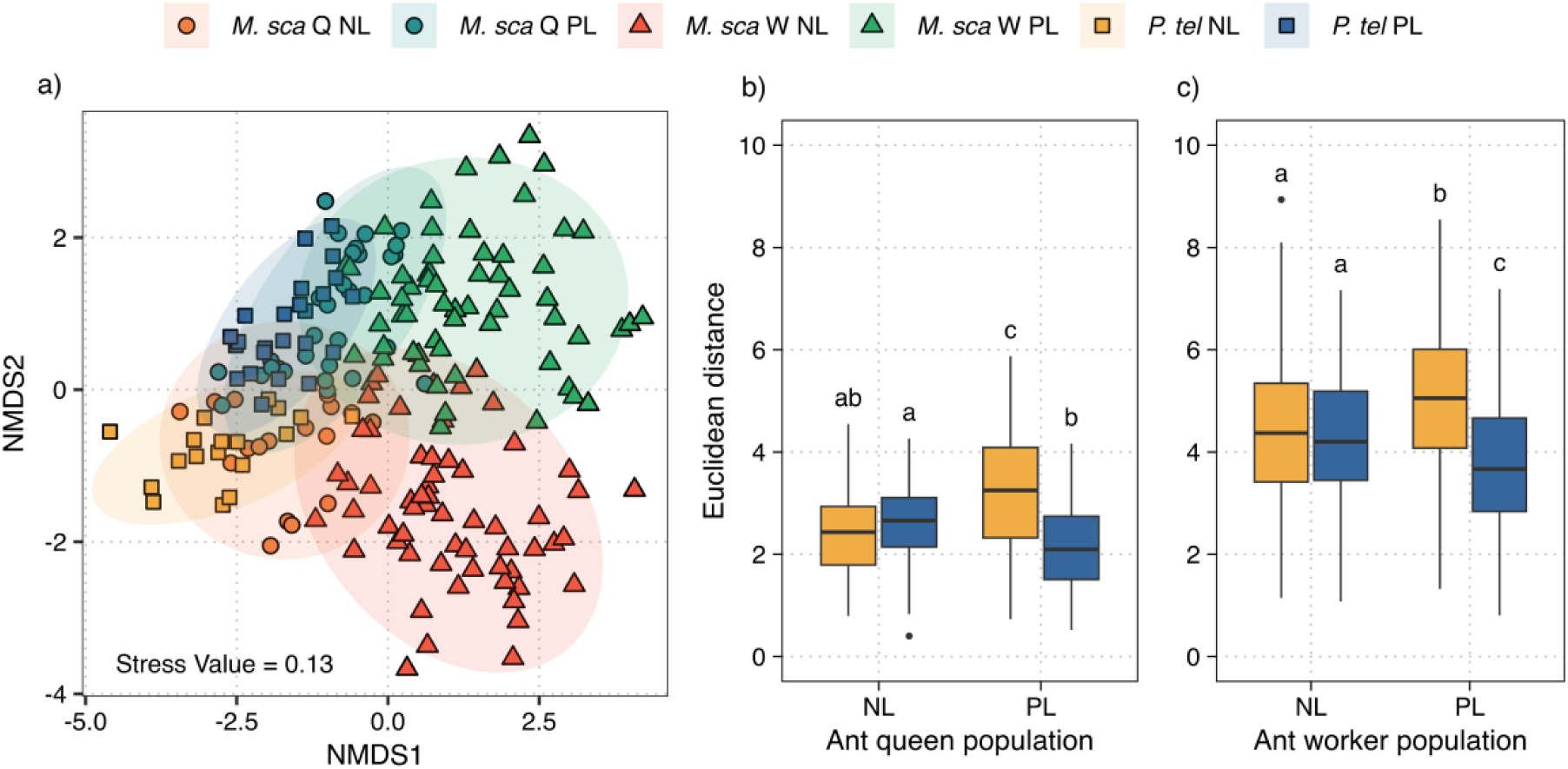
Comparison of vibroacoustic signals emitted by *P. teleius* caterpillars, ant queens and workers based on normalized Euclidean distances calculated using single unit parameters. (a) NMDS ordination. Each point represents ‘average’ values of pulse parameters calculated over a train of pulses (see Material and Methods for more details). Legend abbreviations: *M. sca* (*M. scabrinodis*), *P. tel* (refers to (*P. teleius*), Q (queen), W (worker), NL (the Netherlands), PL (Poland). (b–c) Euclidean distances between: (b) caterpillars and queens, and (c) caterpillars and workers. Horizontal bars represent the median, boxes represent the 25% and 75% quartiles and whiskers the maximum and minimum values. Dots represent outliers. Different letters above boxplots denote pairwise significant differences based on estimated marginal means tests.

The vibroacoustic similarity was also analyzed fitting the Euclidean distances among queen/ant workers and *P. teleius* caterpillars in a generalized linear model. The differences in vibroacoustic signals were significantly affected by the origin of the caterpillars, but not directly influenced by the origin of the host ants (Table S4a and 5a). The reintroduced caterpillars produced a vibroacoustic signal more similar to that of their sympatric host ants. Similarly, Polish caterpillars exhibited a signal profile comparable to that of their sympatric host ants. Overall, the caterpillars from Poland had the closest vibroacoustic signal when compared to their Polish sympatric host ants, but no difference was found between Polish and Dutch caterpillars when they were compared to Dutch host ants (Figure 2b and 2c; Table S4b and 5b).

The vibroacoustic signals emitted by *M. scabrinodis* queens and *P. teleius* caterpillars from Poland share similar values for all the estimated vibroacoustic parameters (Figure S7; Table S6 and 7). Most of the values of the Dutch *P. teleius* caterpillars are similar to those of their sympatric host queens, apart from the root-mean-square values (Figure S7d; Table S7d). Moreover, the parameters related to sound amplitude (the root-mean-square and energy of the peak frequency) differed between *P. teleius* caterpillars from the Dutch and Polish population (Figure S7d and 7e; Table S7d and 7e). The amplitude parameters not only differed in the caterpillars, but also showed a considerable divergence among ant populations.

During the playback experiments, worker ants displayed no aggressive or alarmed behaviors. Instead, we observed five positive responses. Notably, the digging behavior, although observed infrequently, was never triggered by the white noise control stimulus. The results indicated significant differences in ant responses to the five vibroacoustic signals, when considering the total behaviors exhibited per minute of observation (Figure 3 and Table S8). When ants were exposed to caterpillar sounds, the workers of each population reacted significantly more to those produced by their sympatric parasite (Polish workers: z = −5.079, p < 0.001; Dutch workers: z = 4.323, p < 0.001).

**Figure 3.**
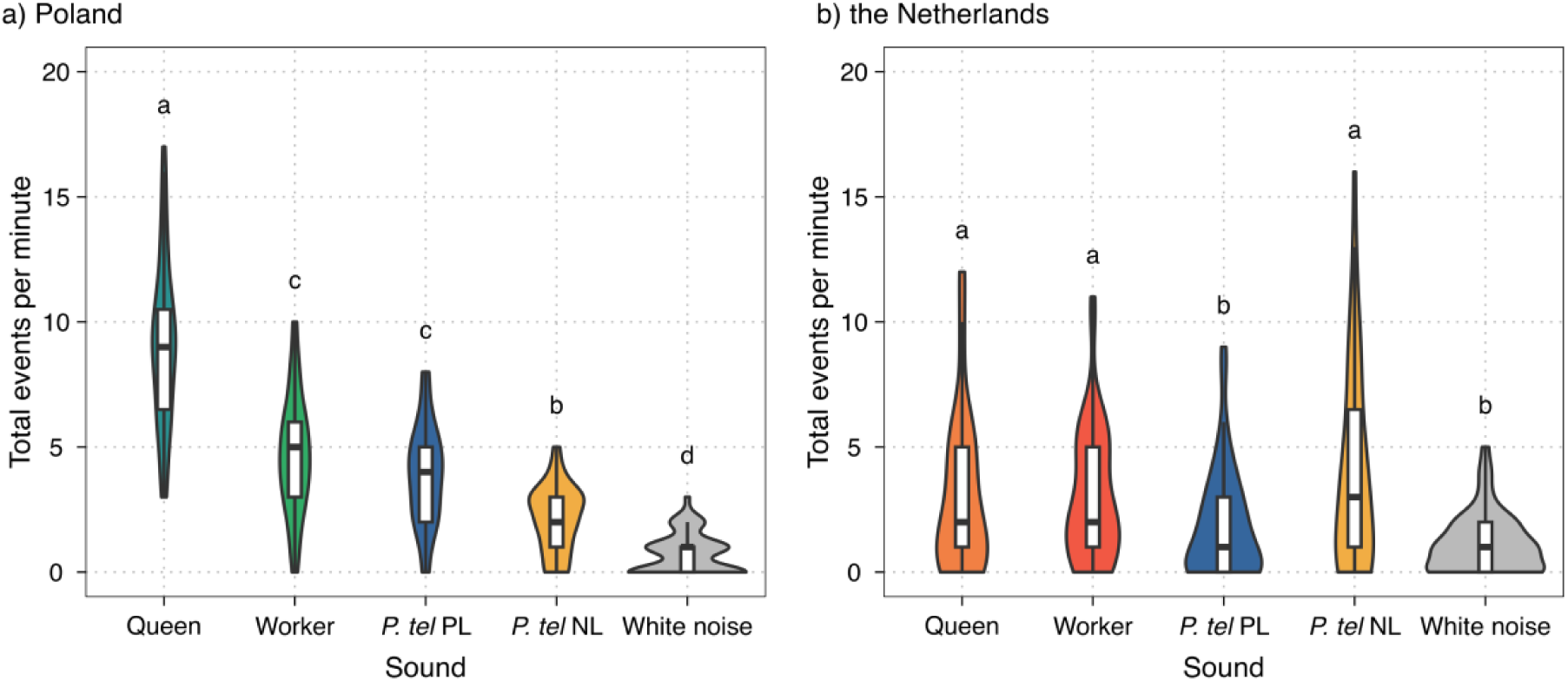
Behavioural responses of *M. scabrinodis* workers from (a) Poland and (b) the Netherlands to vibroacoustic emissions of sympatric *M. scabrinodis* queens and workers, and the emission of sympatric and allopatric *P. teleius* caterpillars and to a control signal (white noise). Horizontal bars represent the median, boxes represent the 25% and 75% quartiles and whiskers the maximum and minimum values. Dots represent outliers. Different letters above boxplots denote pairwise significant differences based on estimated marginal means tests.

### Behavioural responses

During the adoption experiment, the population of the caterpillars influenced the number of antennation events performed by the host ants (Table S9a). However, the number of antennation events did not differ among groups (Figure 4a; Table S9b). Additionally, the population of the ants and the interaction among the population of the caterpillars and ants did not show any significant effect (Table S9a). The number of positive behaviors performed by the Polish host ants to their sympatric caterpillars was significantly higher compared to the positive reaction to the reintroduced caterpillars, but no other significant differences were found (Figure 4b; Table S10b). The population of the caterpillars and ants did not show to be a trigger for a change in the number of positive behaviors (Table S10a). However, the interaction between both variables showed a cross-over effect with a higher median number of positive behaviors for the sympatric combinations. The number of negative behaviors received by *P. teleius* caterpillars from the host ants was significantly lower for the sympatric Polish host-parasite combination compared to any other group (Figure 4c; Table S11b). The caterpillar population did not show to be a general trigger for a change in the number of negative behaviors in any case, but the ant population and the interaction among caterpillar and ant population influenced the number of negative behaviors (Table S11a).

**Figure 4.**
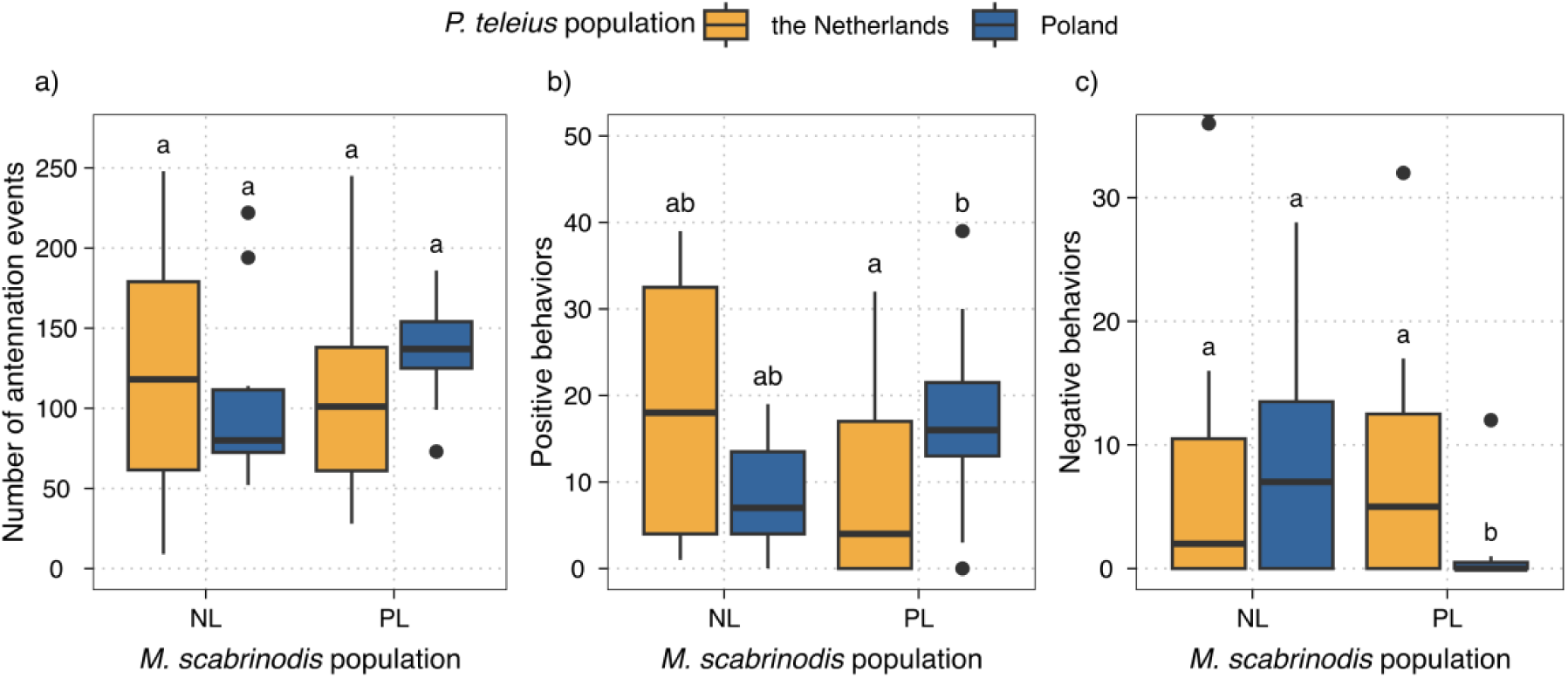
*M. scabrinodis* ant and *P. teleius* caterpillar behavioural cross-population experiment results for (a) number of antennation events, (b) positive behaviours and (c) negative behaviours. The color indicates the population of origin for *P. teleius*: yellow (the Netherlands) and blue (Poland). Horizontal bars represent the median, boxes represent the 25% and 75% quartiles and whiskers the maximum and minimum values. Dots represent outliers. Different letters above boxplots denote pairwise significant differences based on estimated marginal means tests.

None of the studied variables significantly affected the number of drops produced by the caterpillars (Figure S8a; Table S12). However, the number of drops was highly positively correlated with the number of positive behaviors performed by ants towards caterpillars (r^2^ = 0.49, p < 0.001; Figure S8b), but we did not find any correlation between the number of drops and the number of ant negative behaviors (r^2^ = −0.16, p = 0.259; Figure S8c).

*P. teleius* caterpillars from Poland had the highest probability of being adopted among all combinations (Figure 5a). The population of the caterpillars influenced the adoption success, whereas the population of the host ants did not show any significant effect on the proportion of adopted individuals (Table S13). The survival of caterpillars after adoption was significantly affected by the host ant population and marginally influenced by the caterpillar population (Figure 5b; Table S14a). The hazard ratios showed a significant increase of the survival probability of the caterpillars after being adopted by the Polish host ants (HR = 0.49, p = 0.037) and a marginally significant increase of the survival when the caterpillars belonged to the Polish population (HR = 0.53, p = 0.052) (Table S15). The survival probability for the Polish caterpillars with their sympatric host ants showed significantly higher survival probability with respect to the Dutch caterpillar survival probability with their sympatric host ants (Table S14b).

**Figure 5.**
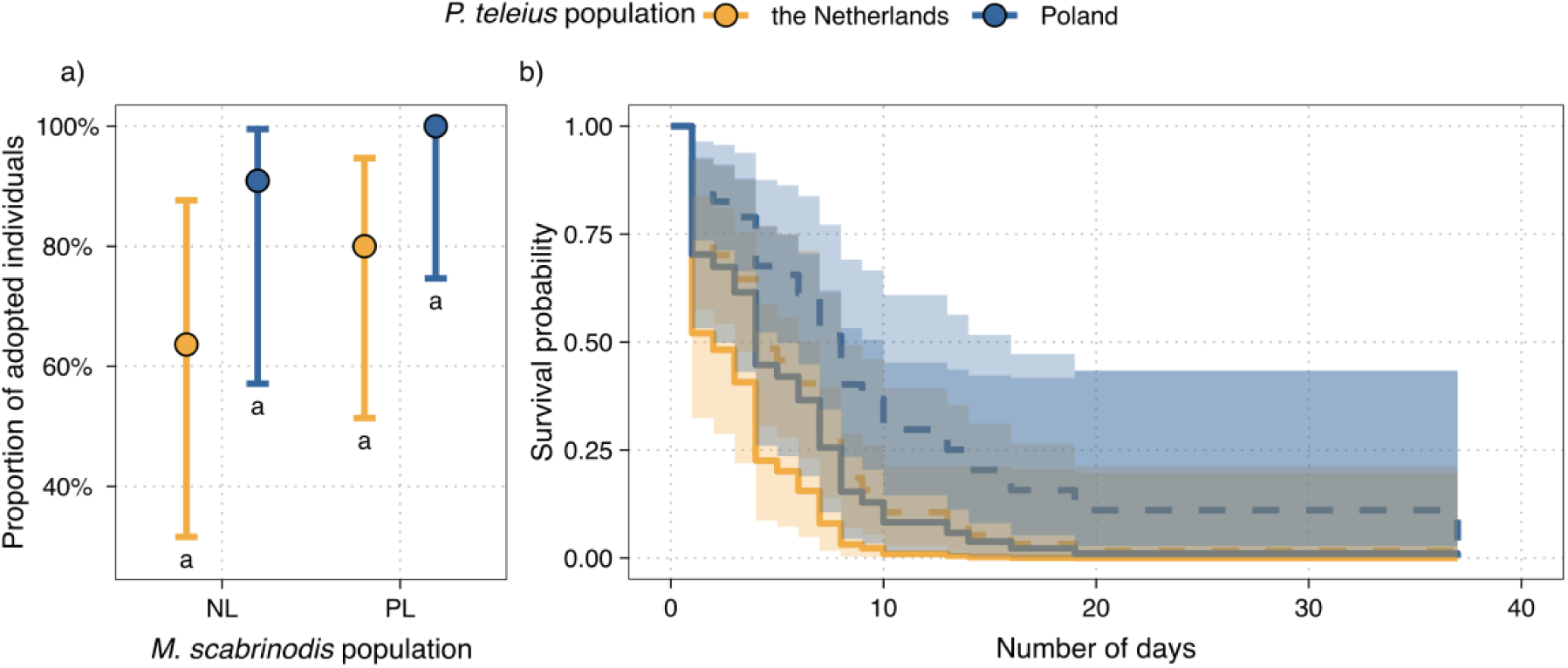
*M. scabrinodis* ant and *P. teleius* caterpillar behavioural cross-population experiment results for (a) adoption success and (b) caterpillar survival probability. The color indicates the population of origin for *P. teleius*: yellow (the Netherlands) and blue (Poland). In plot (a) different letters above boxplots denote pairwise significant differences based on estimated marginal means tests. In plot (b) solid lines refers to the host ant Dutch population, while dotted ones refers to the Polish population.

## Discussion

Chemical and acoustic signals play a vital role in communication within insect societies, and they are often mimicked by social parasites to deceive and exploit their host colonies [38,41,42]. The subversion of these communication signals has been extensively studied in the butterflies of the genus *Phengaris*, which integrate into the colonies of *Myrmica* ants [20,43]. The degree to which the caterpillar can successfully mimic such signals plays a fundamental role in their survival.

### Cuticular hydrocarbon adaptations

We show that thirty years after their reintroduction, descendants of *P. teleius* in the Netherlands have developed different chemical signals from the Polish population they originated from. These differences were observed during the pre-adoption phase, when the profiles of Dutch caterpillars matched their local hosts better than their source host population. High caterpillar mortality rates are recorded during the phases of adoption and initial colony integration [6,44,45], representing critical moments in the life cycle. However, even if these initial phases generate a strong selective pressure on *Phengaris* caterpillars and may have favoured, among the reintroduced individuals, those with chemical characteristics more suited to deceive the colonies of the Dutch host ant, we cannot assume that this alone triggered the change we detected in pre-adoption cuticular hydrocarbons. A large-scale analysis, based on the collection and comparison of data from different European populations, has highlighted that in *P. teleius* the use of the host is regulated by frequency-dependent selection mechanisms, in which parasites tend to exploit the most abundant ant species [22]. If that would be the mechanism explaining our results we would expect a similar pattern in the Polish caterpillars, being more similar to their local host ants. However, our results suggest that the Polish caterpillars are more similar to the Dutch ants. In this context, the most frequently exploited host ants may develop counter-adaptations over time, with an increase in their ability to recognize and repel parasitic caterpillars. Such a dynamic could be observed in Polish ant populations, which have a longer coevolutionary history with *P. teleius* and have been subjected to parasitism pressure for several generations. Beyond these biotic factors, the differences in pre-adoption chemical profiles could also reflect a direct effect of local climate conditions. Mean air temperatures are lower in the Dutch site than in the Polish one [27], and exposure to colder climates is associated with an increase in branched CHCs to improve cuticle fluidity [46,47]. Consistently, Dutch caterpillars present a higher abundance of certain branched compounds such as 3,7,13-trimethyl-C29 and 13,17-dimethyl-C33. In line with this hypothesis, our data show a slight lower similarity between the chemical profiles of pre-adoption caterpillars and host ants in Poland than in the Netherlands. This scenario can be interpreted as evidence of environmental constrains and, mainly, coevolutionary dynamics between *Phengaris* butterflies and *Myrmica* ants, as already demonstrated for *P. teleius* with *M. scabrinodis* [48] and for *P. alcon* with *M. rubra* [10]. Thus, the higher degree of chemical similarity of the reintroduced parasite observed during pre-adoption stage could have also been facilitated by the absence of parasitic pressure experienced by Dutch ants during the fourteen years following the extinction of *P. teleius* in the Netherlands until its reintroduction [26].

In the post-adoption phase, Polish *P. teleius* caterpillars show a greater chemical similarity with its sympatric host ants, suggesting a more effective integration due to a longer coevolutionary history. Although chemical mimicry within the ant colony in *P. teleius* is less sophisticated than that observed in cuckoo species of the genus *Phengaris* [8,9]. Our results suggest that even if less efficiently than cuckoo species, *P. teleius* caterpillars are able to adapt chemically to their host ants after adoption. Additionally, the differences in CHC compounds between caterpillars of the two populations are smaller after adoption than during the pre-adoption phase. This is likely due to the facilitation by the stable microclimatic conditions inside the ant nest and the chemical changes triggered by adoption, such as the detected increase of 3-methyl-C23. This compound is critical for recognizing intruders within the colony [49,50]. Specifically, the Dutch caterpillars had a significantly higher abundance of 3-methyl-C23 than Polish caterpillars before the adoption, but this difference disappeared after adoption during the CHC profiles changes. This variation could reflect a specific adaptation mechanism to the host colony in the less coevolved conditions of the reintroduced population, suggesting a possible active role of this compound in chemical integration.

### Vibroacoustic signal adaptations

Similarly to chemical signals, we found that the reintroduced caterpillars produce vibroacoustic signals that are different from those emitted by Polish caterpillars. In particular, these signals are more similar to the stridulations of Dutch ants than to Polish ones, indicating a process of local adaptation to the new host. In addition, vibroacoustic signals emitted by caterpillars are generally more similar to those of the queen than to those of workers. But unlike the chemical adaptations, vibroacoustic emissions of pre-adoption stage showed a greater proximity between caterpillars and their sympatric host ants, despite the reintroduced caterpillar not showing a closer signal with their local hosts than the Polish caterpillar, which never coevolved with the Dutch ants before. However, it should be noticed that the level of similarity between Dutch and Polish caterpillars to the ants is almost identical, and in the playback experiment the ants reacted more positively towards vibroacoustic signals produced by their sympatric caterpillars, giving evidence of local adaptation and indicating a possible main role of vibroacoustic signals during the initial adoption phase. The effectiveness of acoustic mimicry in *Phengaris* varies depending on the developmental stage of the butterfly caterpillar (pre- and post-adoption) and its trophic strategy (cuckoo vs. predatory) [12]. In the post-adoption stage, vibroacoustic signals emitted by cuckoo species elicit stronger responses from workers than those produced by predatory species, thus allowing the cuckoo parasite to achieve a high status with preferential feeding and protection from the host ants. On the contrary, in the pre-adoption stage, the calls of predatory species attract the attention of foragers more than the signals emitted by cuckoo species, presumably compensating for their less accurate chemical mimicry and facilitating retrieval by ants [12]. Our results suggest that acoustic signals play a primary role in the pre-adoption phase, not only attracting the ant foragers, but also directly contributing to the process of specific recognition.

Furthermore, it is not only the general similarity of sound that is relevant; also the values of specific parameters of vibroacoustic signals could be determinants of sound mimicry in *Phengaris* butterflies. For instance, the moth *Achroia grisella* is able to distinguish sounds that differ in only one component and shows a preference for more regular and higher amplitude signals [51]. Amplitude parameters (i.e. peak frequency energy) differed between *Phengaris* and *Myrmica* populations, but signals were still significantly similar between caterpillars and sympatric queens. Sound intensity has been identified in *Phengaris*–*Myrmica* systems as one of the key parameters in the similarity between caterpillar and queen signals [12] and our results support the idea that intensity may represent a crucial component of the signals that drive differentiation between populations and enhance vibroacoustic deception towards *Myrmica* hosts.

### Behavioural responses

The behaviours observed between ants and caterpillars during the adoption process can be used as an indirect indicator of the effectiveness of chemical and acoustic strategies used by *P. teleius* to be adopted and retrieved into the ant colony. Our study showed a clear pattern, with the lowest number of negative behaviours performed by Polish ants towards their sympatric parasite caterpillars. Although not statistically significant, also ants from the Netherlands showed a lower incidence of negative behaviours towards their sympatric caterpillars. Antennations and positive behaviours, although not significantly different among groups, were observed more frequently in both sympatric host-parasite combinations. Antennations, in particular, could play an important role in the initial phases of recognition, allowing workers to perceive cuticular compounds through their antennae and compare the detected chemical profile with the internal reference template [13].

A previous adoption experiment comparing sympatric and allopatric *P. teleius* caterpillar populations found a higher number of positive behaviours in the sympatric group, but no difference in antennations and negative behaviours [44]. However, positive behaviours are difficult to interpret as they could be bias. In our study, we observed a strong correlation between the number of sugar drops and positive behaviours, suggesting that less adapted caterpillars may use sugar secretions as a compensatory strategy, thus reducing the differences observed between sympatric and allopatric combinations. Conversely, the number of negative behaviours represents a reliable metric to assess the success of integration, as we found no correlation between the number of drops and aggressive responses. This suggests that sugar secretion is not sufficient to prevent aggression when caterpillars are recognized as intruders.

We found no statistically significant differences in the proportion of adopted caterpillars between the reintroduced and source populations, which is consistent with the chemical results obtained for in the pre-adoption phase, indicating a comparable level of mimicry between Polish and Dutch caterpillars at this stage of development. Despite the absence of statistical differences, it is important to highlight that caterpillars from Poland showed the highest probability of being adopted by their sympatric hosts. This could reflect the more similar vibroacoustic signals found in the Polish caterpillars to their host ants and a higher level of adaptation resulting from a longer coevolutionary history. Similar results, where sympatric populations show a higher probability of adoption, have already been observed in *Phengaris nausithous* and *P. teleius* [48,52]. However, it has been reported that in *P. alcon* more successful adoption can also occur in the presence of allopatric ants [53].

Furthermore, Polish caterpillars, in the presence of their host ants, showed a higher probability of survival than the reintroduced Dutch group. This result is in line with post-adoption chemical data, which indicate that Polish caterpillars are able to better adapt to sympatric colonies, thus increasing their chances of survival within the host colony.

## Conclusions

Thirty generations after the reintroduction of *P. teleius* in the Netherlands were sufficient to reveal differences in the adaptations of the reintroduced caterpillars: both chemical and acoustic signals were different from those of the caterpillars of the Polish population of origin. The shorter generational cycle of *P. teleius* may allow the parasitic butterfly to evolve more rapidly than its host ants. As a result, the butterfly may have a relative advantage in adapting to the parasitic relationship, potentially outpasing the speed at which ants develop defenses against parasitism. This difference in generational timing could significantly influence coevolutionary dynamics, affecting the pace of local adaptations in both the butterfly and the ants.

Our results suggest that the two communication channels used by the caterpillars to deceive the host ants – the chemical and the acoustic one – follow different coevolutionary trajectories. As suggested by [12], cuckoo and predatory caterpillars of the genus *Phengaris* may invest differently in chemical and acoustic signals, also considering the different needs in the pre- and post-adoption phases. In predatory species, which rely on long adoption rituals, caterpillars in the pre-adoption phase may rely more on acoustic emissions to support chemical mimicry. It could be due to differences in the evolutionary costs associated with the two communication channels: acoustic mimicry, in theory, could be more easily modulated, while chemical mimicry faces more constraints during the pre-adoption phase. Chemical mimicry may depend on multiple factors before adoption, such as the number of host species, ant counteradaptations, or environmental conditions; consequently, its plasticity and adaptability may only increase after adoption, where it could be less constrained by these initial hurdles.

We provide evidence that *P. teleius*, considered the most generalist species among all butterflies of the genus towards its host ants [44,54–56], is able to evolve novel adaptations or show high phenotypic plasticity, adapting to new local populations of its host ants. The levels of chemical and acoustic mimicry detected in reintroduced caterpillars, although they do not reach the levels of effectiveness observed in the source population, appear to be sufficient to ensure adoption by host ants and have proven to be adequate to allow the survival and growth of a currently viable *P. teleius* population at the reintroduction site.

## Acknowledgements

The study was funded by the Polish National Science Centre (NCN) grant 2018/31/B/NZ8/03476. National State Forestry, Natuurmonumenten and the Province of Northern Brabant gave us permission to access the nature reserve and carry out the survey. Permission for butterfly capture in Kraków was given by the Regional Directorate for Environmental Protection in Kraków (decisions DZP-WG.6401.01.29.2019.ep.eb). We would like to thank Gema Trigos Peral for her help during the fieldwork in Poland, Andrea Zagato for carrying out sound annotation and performing part of the playback observations and Jerzy Romanowski for allowing us to use their microbalance.

## CRediT authorship contribution statement

Daniel Sánchez-García: Conceptualization, Formal analysis, Investigation, Data Curation, Writing - Original Draft, Visualization. Irma Wynhoff: Conceptualization, Methodology, Investigation. Patricia d’Ettorre: Investigation. Chloé Leroy: Investigation. Joanna Kajzer-Bonk: Investigation. István Maák: Investigation. Francesca Barbero: Investigation, Writing - Review & Editing. Luca Pietro Casacci: Conceptualization, Methodology, Formal analysis, Investigation, Writing - Review & Editing. Magdalena Witek: Conceptualization, Methodology, Investigation, Writing - Original Draft, Supervision, Project administration, Funding acquisition.

## Conflict of Interest Statement

The authors declare no conflict of interests.

## Notes

https://doi.org/10.5281/zenodo.17791047

### Competing Interest Statement

The authors have declared no competing interest.

