## Supplementary Figures and Tables for "Ongoing coevolution between reintroduced *Phengaris teleius* butterflies and their *Myrmica* host ants"

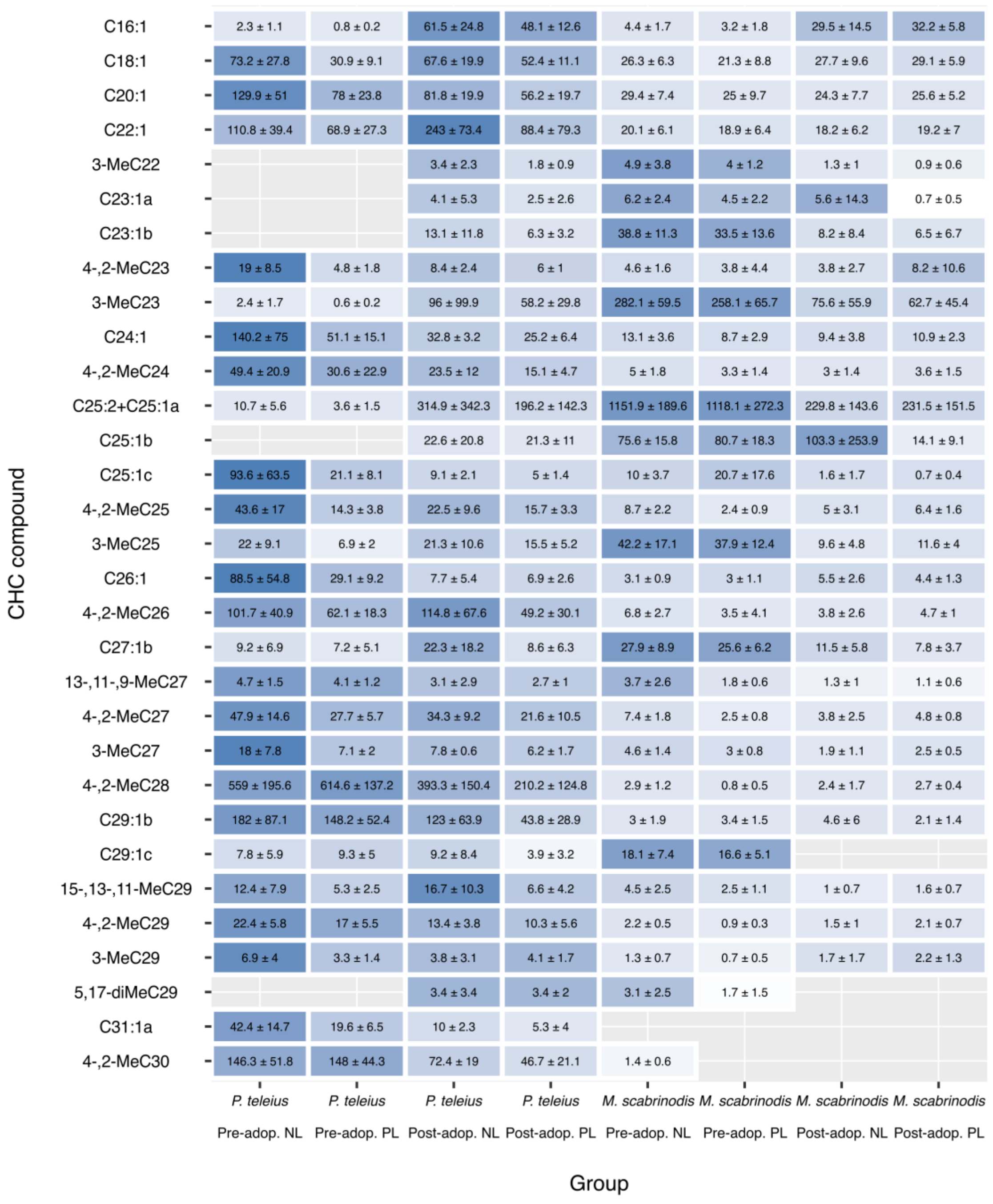

Figure S1. Graphical representation of the CHC compounds analyzed in the study. Blue tiles indicate the presence of the CHC in the different groups. The values inside the tiles indicate the absolute abundance of the CHCs (ng) per mg of sample dry mass and their standard deviation.

Supplementary material. Sánchez-García, D. et al. Ongoing coevolution between reintroduced *Phengaris teleius* butterflies and their *Myrmica* host ants.

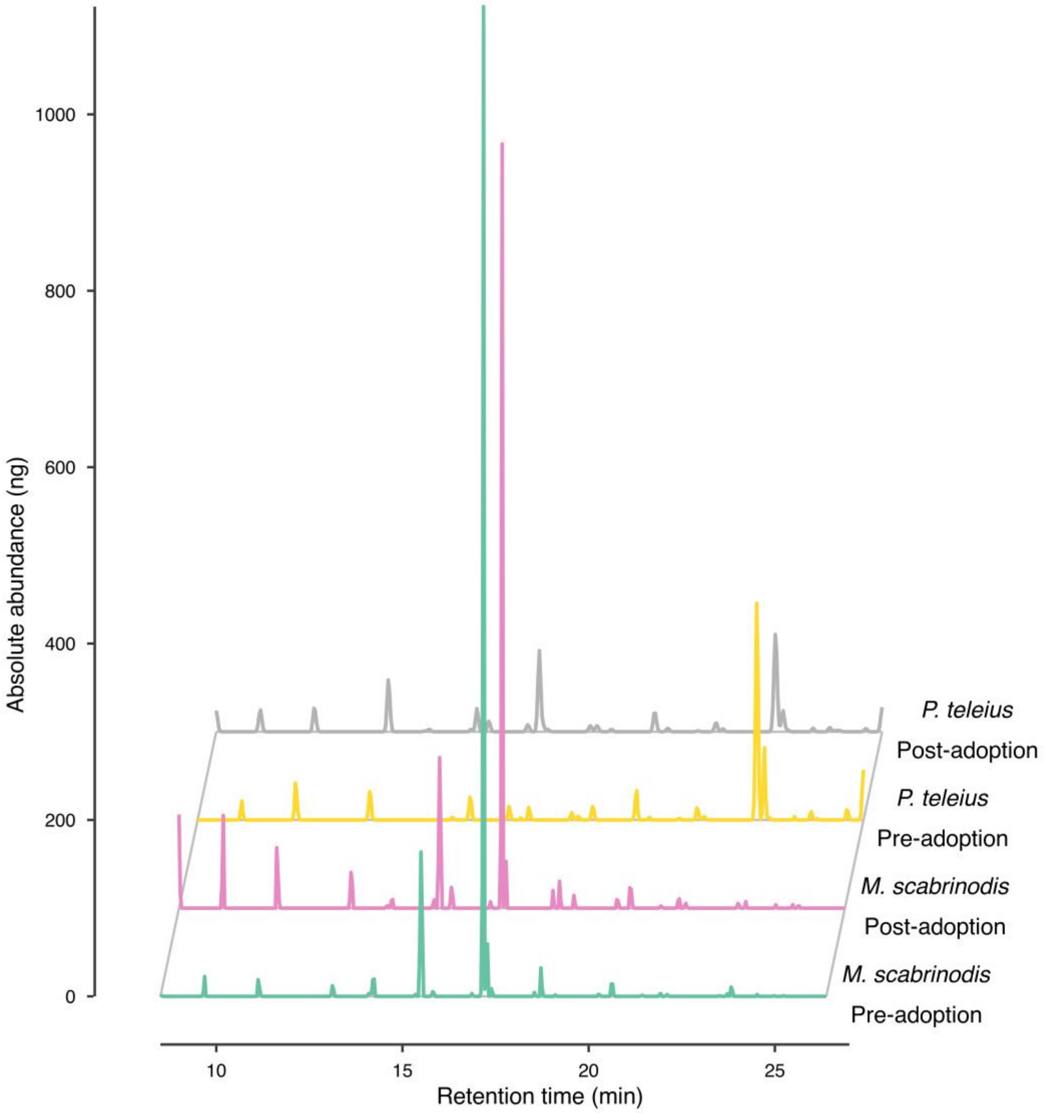


Figure S2. Chromatogram representation of the CHC profile from the different studied groups. The X axis indicates the retention time in which the compound was detected by the GC-MS analysis. The Y axis indicates the absolute abundance (ng) of the CHC compounds per mg of sample dry mass.

Supplementary material. Sánchez-García, D. et al. Ongoing coevolution between reintroduced *Phengaris teleius* butterflies and their *Myrmica* host ants.

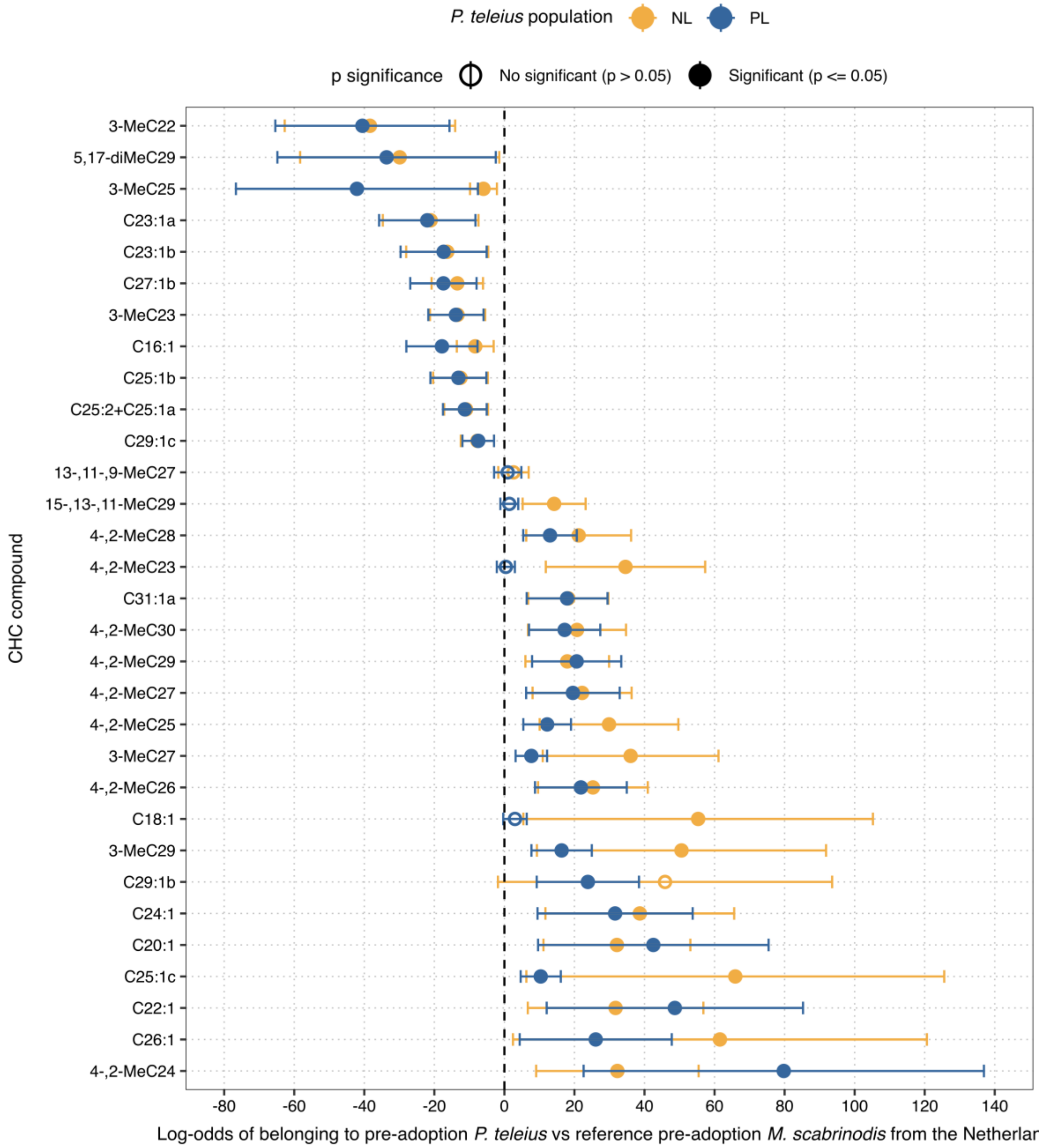


Figure S3. Forest plot of log-odds estimates of cuticular hydrocarbon (CHC) compounds predicting membership in pre-adoption *P. teleius* populations (the Netherlands and Poland) versus the reference pre-adoption *M. scabrinodis* population from the

Netherlands. Dots represent GLM estimates for individual compounds, with error bars showing 95% confidence intervals. Filled circles indicate significant compounds ($p \leq 0.05$) and open circles indicate non-significant compounds. Colors indicate the population of *P. teleius*, with NL (yellow) referring to Netherlands and PL referring to Poland (blue). Compounds are ordered by decreasing log-odds estimate.

Supplementary material. Sánchez-García, D. et al. Ongoing coevolution between reintroduced *Phengaris teleius* butterflies and their *Myrmica* host ants.

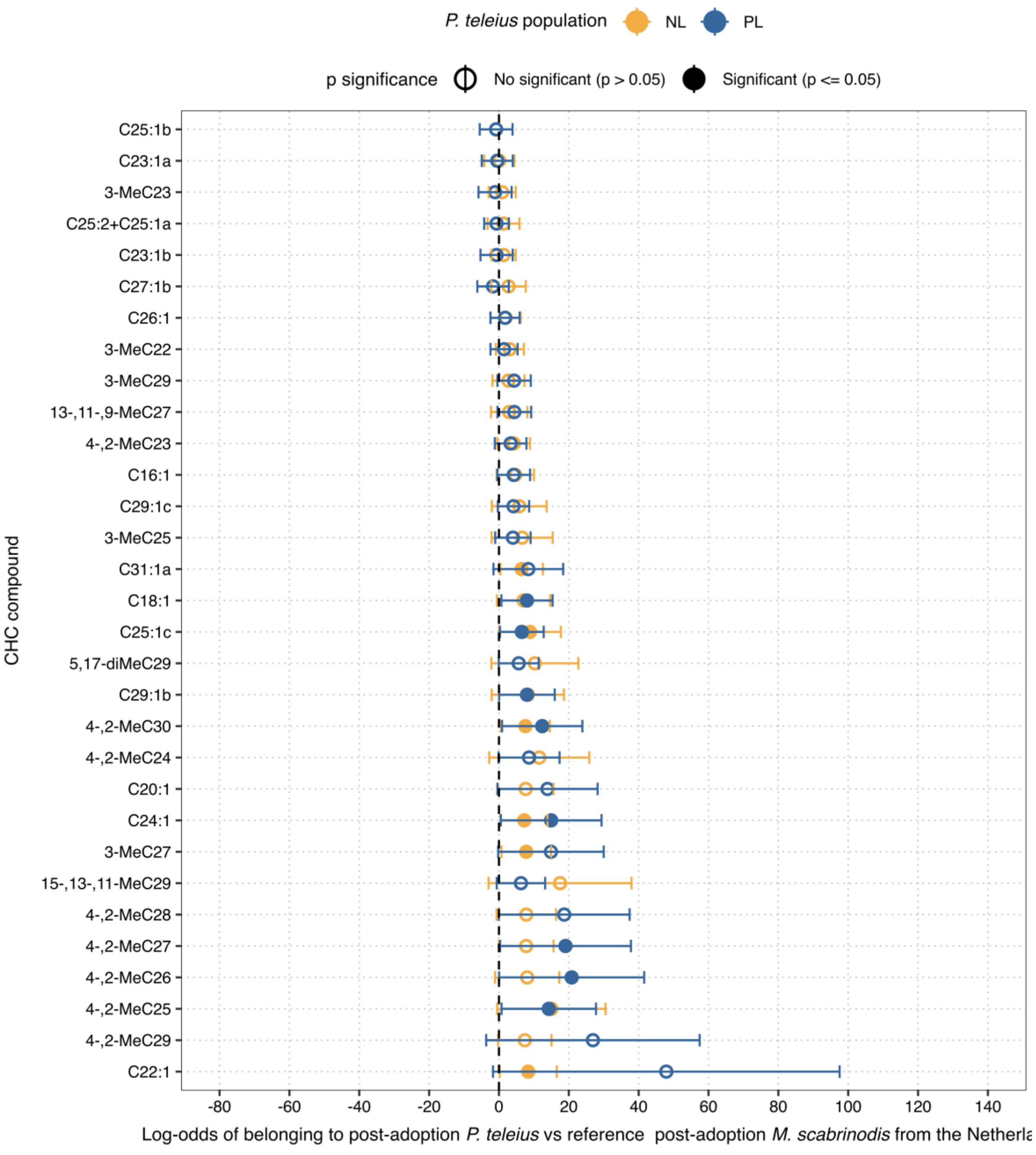


Figure S4. Forest plot of log-odds estimates of cuticular hydrocarbon (CHC) compounds predicting membership in post-adoption *P. teleius* populations (the Netherlands and Poland) versus the reference post-adoption *M. scabrinodis* population from the

Netherlands. Dots represent GLM estimates for individual compounds, with error bars showing 95% confidence intervals. Filled circles indicate significant compounds ($p \leq 0.05$) and open circles indicate non-significant compounds. Colors indicate the population of *P. teleius*, with NL (yellow) referring to Netherlands and PL referring to Poland (blue). Compounds are ordered by decreasing log-odds estimate.

Supplementary material. Sánchez-García, D. et al. Ongoing coevolution between reintroduced *Phengaris teleius* butterflies and their *Myrmica* host ants.

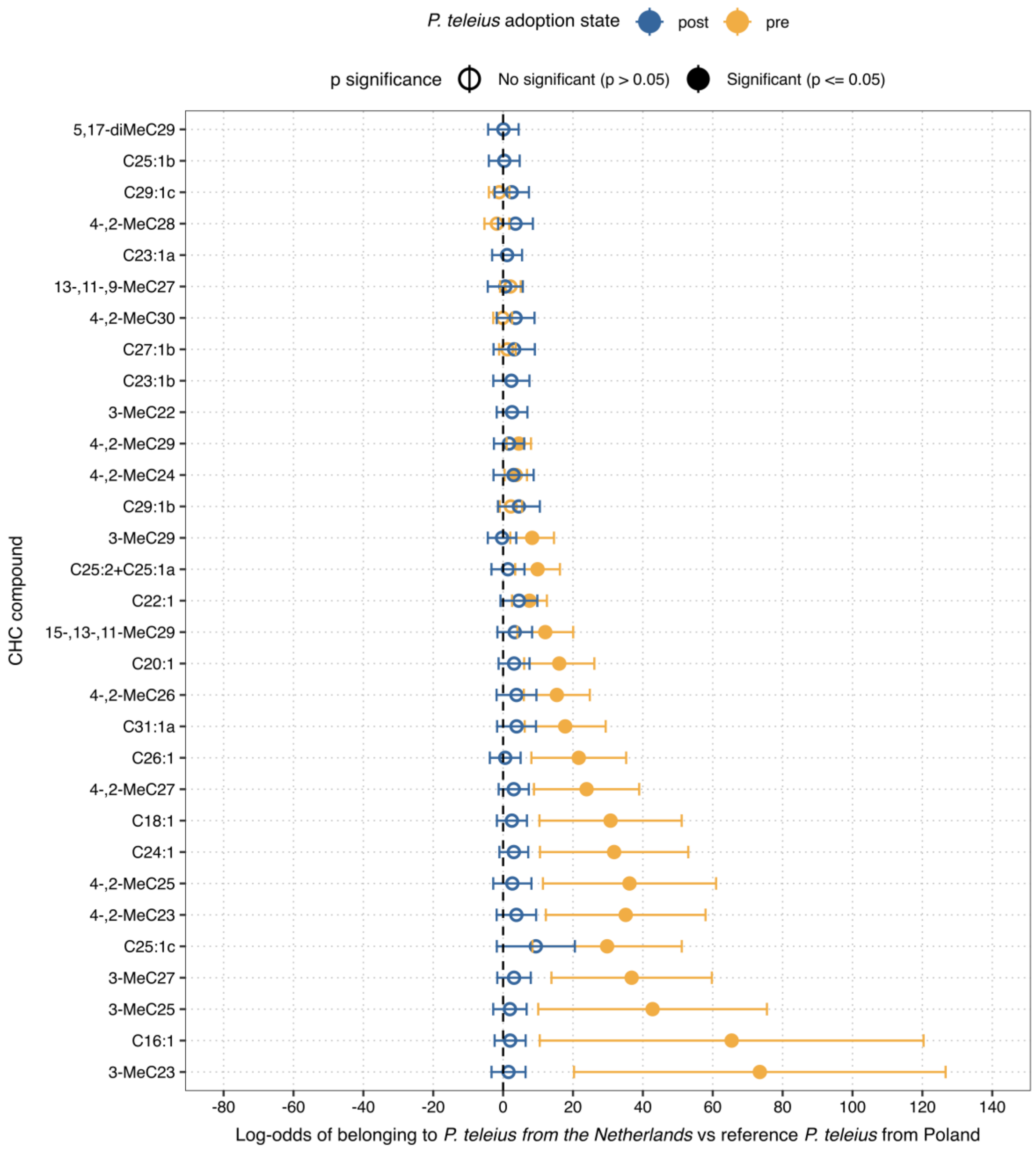


Figure S5. Forest plot of log-odds estimates of cuticular hydrocarbon (CHC) compounds predicting membership in *P. teleius* from the Netherlands (the Netherlands and Poland) versus the reference *P. teleius* from Poland for the pre- and post-adoption state.

Dots represent GLM estimates for individual compounds, with error bars showing 95% confidence intervals. Filled circles indicate significant compounds ($p \leq 0.05$) and open circles indicate non-significant compounds. Colors indicate the adoption state of *P. teleius*, with pre (yellow) referring to pre-adoption and post referring to post-adoption (blue). Compounds are ordered by decreasing log-odds estimate.

Supplementary material. Sánchez-García, D. et al. Ongoing coevolution between reintroduced *Phengaris teleius* butterflies and their *Myrmica* host ants.

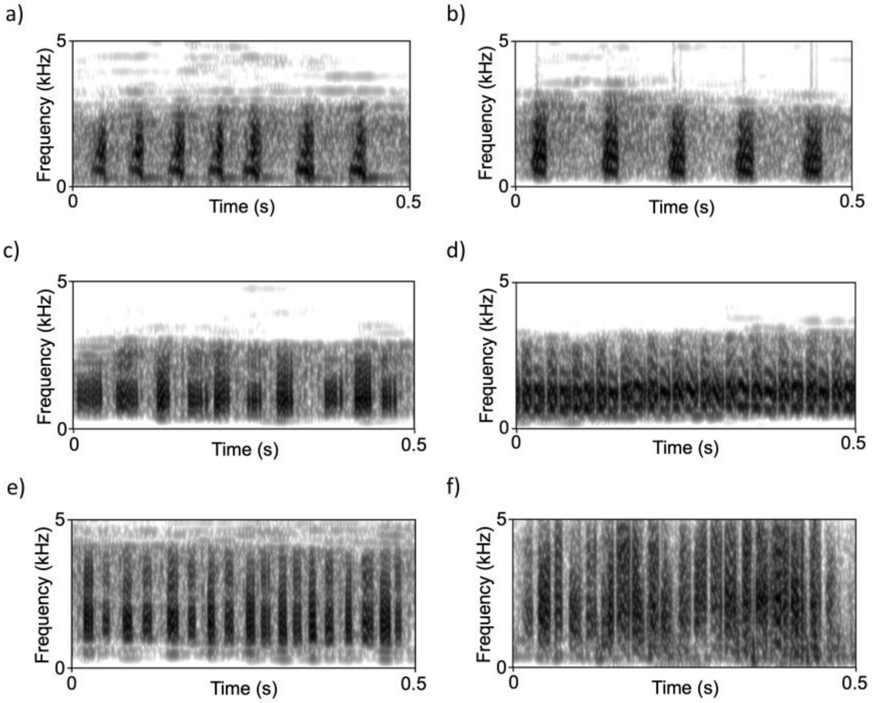


Figure S6. Oscillograms and spectrograms of the stridulations emitted by pre-adoption caterpillars of *P. teleius*, *M. scabrinodis* queens and workers from a), c), e) Poland and b), d), f) the Netherlands. Spectrograms were generated in Praat using the following parameters: window shape = Gaussian, window length = 0.025 s, number of time steps = 1000, number of frequency steps = 500, dynamic range = 70 dB.

Supplementary material. Sánchez-García, D. et al. Ongoing coevolution between reintroduced *Phengaris teleius* butterflies and their *Myrmica* host ants.

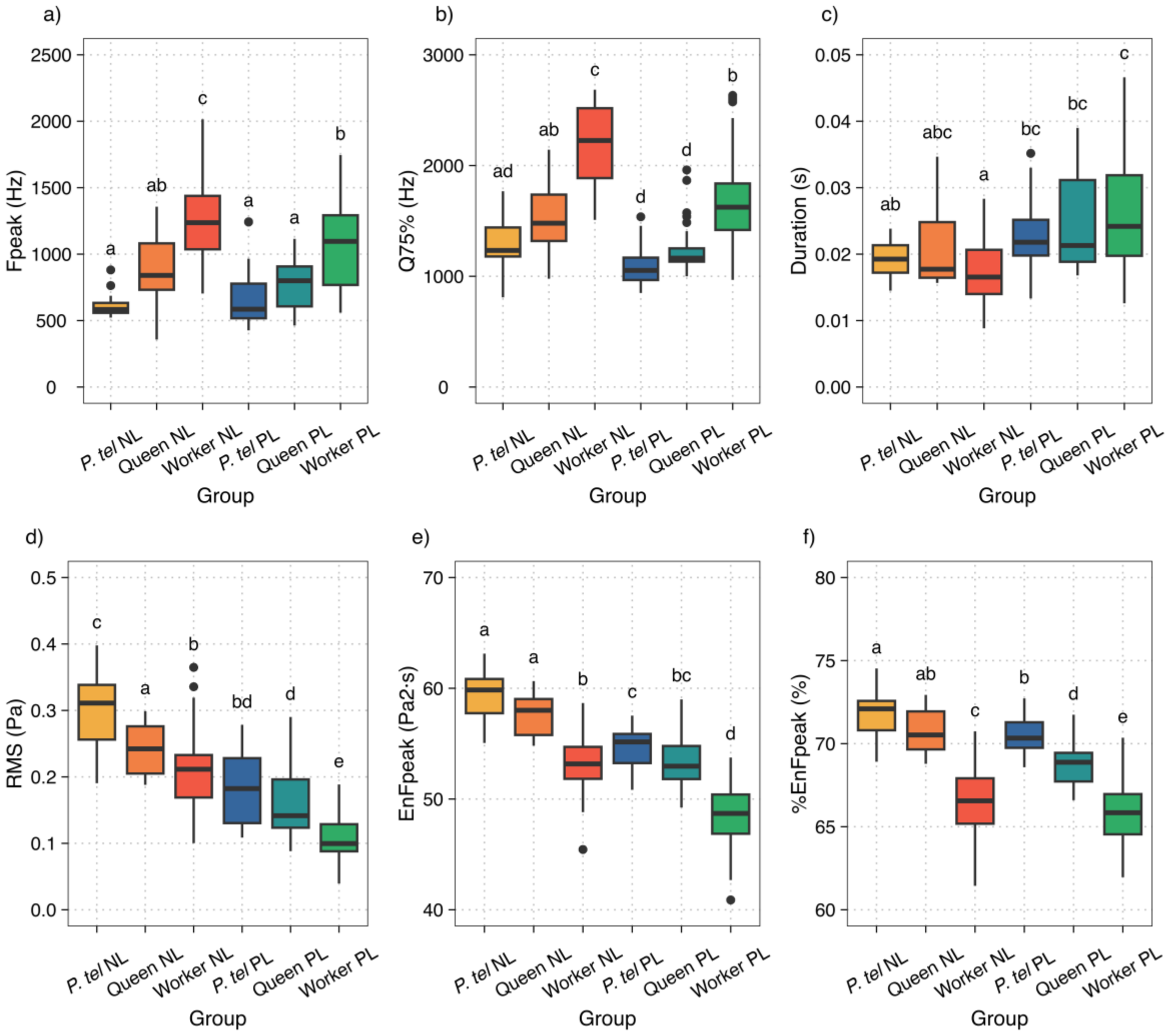


Figure S7. Boxplots of vibroacoustic parameters, i.e., a) frequency peak (Fpeak, Hz), b) third frequency quartile (Q75%, Hz), c) unit duration (Duration, s), d) root-mean-square (RMS, Pa), e) energy of the frequency peak (EnFpeak, $Pa^2 \cdot s$), f) percentage of the energy of the frequency peak over the total energy (%EnFpeak, %), of stridulation units of the signals emitted by *M. scabrinodis* queens, workers and pre-adoption caterpillars of *P. teleius* from the Polish and Dutch metapopulations. Horizontal lines represent median values, the boxes the first and third quartiles and whiskers the maximum and minimum values. Dots represent outliers. Lower-case letters above boxplots indicate pairwise significant differences between castes based on an estimated marginal means (EMMs) test.

Supplementary material. Sánchez-García, D. et al. Ongoing coevolution between reintroduced *Phengaris teleius* butterflies and their *Myrmica* host ants.

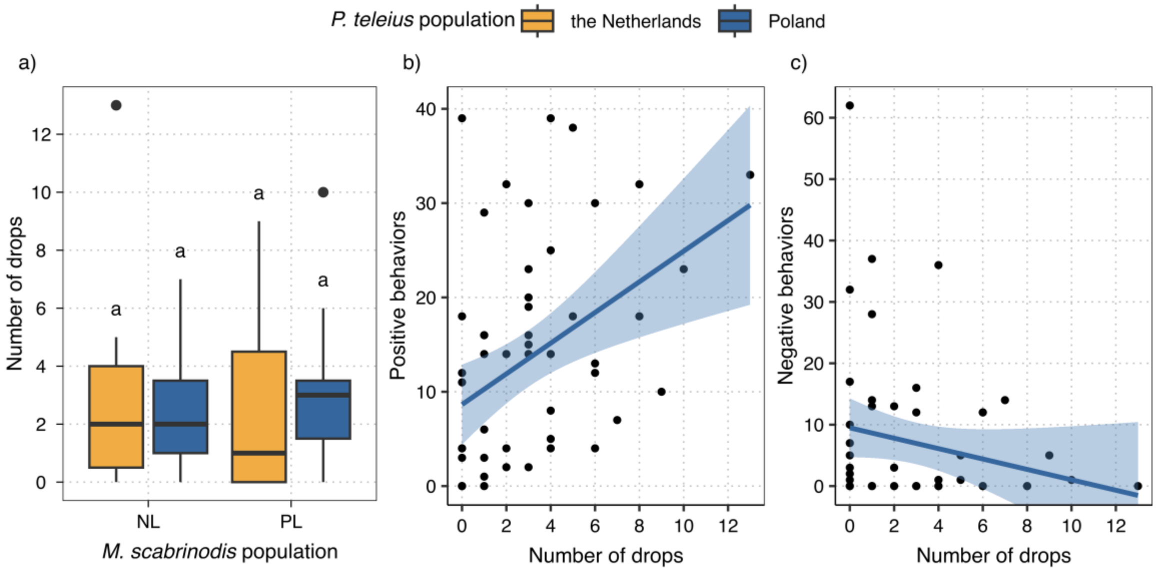


Figure S8. *M. scabrinodis* ant and *P. teleius* caterpillar behavioral cross-metapopulation experiment results: a) number of drops produced by *P. teleius* caterpillars in the different host-parasite combinations. The color of the boxplots indicates the population of origin for *P. teleius*: yellow (the Netherlands) and blue (Poland). Horizontal lines represent median values, the boxes the first and third quartiles and whiskers the maximum and minimum values. Dots represent outliers. Lower-case letters above boxplots indicate pairwise significant differences between groups based on an estimated marginal means (EMMs) test; b) correlation between the number of drops and ant positive behaviors; and c) correlation between the number of drops and ant negative behaviors. The light blue surface represents the 95% interval of confidence.

Supplementary material. Sánchez-García, D. et al. Ongoing coevolution between reintroduced *Phengaris teleius* butterflies and their *Myrmica* host ants.

Table S1. Statistical results from the analysis of pre-adoption cuticular hydrocarbon profile Bray-Curtis distances between *P. teleius* caterpillars and *M. scabrinodis* ants from different populations. a) Generalized linear model (GLM) variable significance. b) Estimated marginal means (EMMs) pairwise comparisons among groups. The first term of each group in the contrast indicates the ant host population; the second term indicates the population of *P. teleius*. NL refers to the Netherlands and PL refers to Poland.

a)

| Variable | d.f. | LR Chisq | p |
|---|---|---|---|
| Ant population | 1 | 9.80 | 0.002** |
| Caterpillar population | 1 | 0.57 | 0.449 |
| Ant population x Caterpillar population | 1 | 98.26 | 0.001*** |

*p ≤ 0.05, **p ≤ 0.01, ***p ≤ 0.001

b)

| Contrast | z | p |
|---|---|---|
| NL-NL vs PL-NL | -2.76 | 0.023* |
| NL-NL vs NL-PL | -0.22 | 0.825 |
| NL-NL vs PL-PL | -3.07 | 0.011* |
| PL-NL vs NL-PL | 2.29 | 0.066 |
| PL-NL vs PL-PL | -1.35 | 0.356 |
| NL-PL vs PL-PL | -3.41 | 0.004** |

*p ≤ 0.05, **p ≤ 0.01, ***p ≤ 0.001

p was adjusted for multiple comparisons

b)

| Contrast | z | p |
| --- | --- | --- |

using the Holm correction

Supplementary material. Sánchez-García, D. et al. Ongoing coevolution between reintroduced *Phengaris teleius* butterflies and their *Myrmica* host ants.

Table S2. Statistical results from the analysis of post-adoption cuticular hydrocarbon profile Bray-Curtis distances between *P. teleius* caterpillars and *M. scabrinodis* ants from different populations. a) Generalized linear model (GLM) variable significance. b) Estimated marginal means (EMMs) pairwise comparisons among groups. The first term of each group in the contrast indicates the ant host population; the second term indicates the metapopulation of *P. teleius*. NL refers to the Netherlands and PL refers to Poland.

a)

| Variable | d.f. | LR Chisq | p |
|---|---|---|---|
| Ant population | 1 | 0.70 | 0.403 |
| Caterpillar population | 1 | 10.61 | 0.001** |
| Ant population x caterpillar population | 1 | 1.25 | 0.263 |

*p ≤ 0.05, **p ≤ 0.01, ***p ≤ 0.001

b)

| Contrast | z | p |
|---|---|---|
| NL-NL vs PL-NL | 0.44 | 0.657 |
| NL-NL vs NL-PL | 2.99 | 0.011* |
| NL-NL vs PL-PL | 3.17 | 0.008** |
| PL-NL vs NL-PL | 2.16 | 0.091 |
| PL-NL vs PL-PL | 3.44 | 0.003** |
| NL-PL vs PL-PL | 1.16 | 0.489 |

b)

| Contrast | z | p |
|---|---|---|

*p ≤ 0.05, **p ≤ 0.01, ***p ≤ 0.001

p was adjusted for multiple comparisons using the Holm correction

110

Supplementary material. Sánchez-García, D. et al. Ongoing coevolution between reintroduced *Phengaris teleius* butterflies and their *Myrmica* host ants.

Table S3. Statistical results from the analysis of pre- and post-adoption cuticular hydrocarbon profile Bray-Curtis distances between *P. teleius* caterpillars and *M. scabrinodis* ants from different populations. a) Generalized linear model (GLM) variable significance. b) Estimated marginal means (EMMs) pairwise comparisons among groups. The first term of each group in the contrast indicates the ant host population; the second term indicates the metapopulation of *P. teleius*. NL refers to the Netherlands and PL refers to Poland.

a)

| Variable | d.f. | LR Chisq | p |
|---|---|---|---|
| Caterpillar population | 1 | 4.55 | 0.033* |
| Ant population | 1 | 6.11 | 0.013* |
| Caterpillar population x ant population | 1 | 0.04 | 0.843 |

*p ≤ 0.05, **p ≤ 0.01, ***p ≤ 0.001

b)

| Contrast | z | p |
|---|---|---|
| NL-NL vs PL-NL | -1.68 | 0.366 |
| NL-NL vs NL-PL | -1.71 | 0.366 |
| NL-NL vs PL-PL | -3.24 | 0.007** |
| PL-NL vs NL-PL | -0.16 | 0.869 |
| PL-NL vs PL-PL | -1.79 | 0.366 |
| NL-PL vs PL-PL | -1.40 | 0.366 |

b)

| Contrast | z | p |
|---|---|---|

*p ≤ 0.05, **p ≤ 0.01, ***p ≤ 0.001

p was adjusted for multiple comparisons using the Holm correction

Supplementary material. Sánchez-García, D. et al. Ongoing coevolution between reintroduced *Phengaris teleius* butterflies and their *Myrmica* host ants.

Table S4. Statistical results from the analysis of vibroacoustic signal Bray-Curtis distances between pre-adoption *P. teleius* caterpillars and *M. scabrinodis* ant queens from different populations. a) Generalized linear model (GLM) variable significance. b) Estimated marginal means (EMMs) pairwise comparisons among groups. The first term of each group in the contrast indicates the ant host population; the second term indicates the population of *P. teleius*. NL refers to the Netherlands and PL refers to Poland.

a)

| Variable | d.f. | LR Chisq | p |
|---|---|---|---|
| Queen population | 1 | 0.18 | 0.675 |
| Caterpillar population | 1 | 23.62 | 0.001*** |
| Queen population x caterpillar population | 1 | 516.45 | 0.001*** |

*p ≤ 0.05, **p ≤ 0.01, ***p ≤ 0.001

b)

| Contrast | z | p |
|---|---|---|
| NL-NL vs PL-NL | -4.77 | 0.001*** |
| NL-NL vs NL-PL | -1.26 | 0.280 |
| NL-NL vs PL-PL | 1.48 | 0.280 |
| PL-NL vs NL-PL | 3.01 | 0.010* |
| PL-NL vs PL-PL | 8.45 | 0.001*** |
| NL-PL vs PL-PL | 2.88 | 0.012* |

*p ≤ 0.05, **p ≤ 0.01, ***p ≤ 0.001

Supplementary material. Sánchez-García, D. et al. Ongoing coevolution between reintroduced *Phengaris teleius* butterflies and their *Myrmica* host ants.

Table S5. Statistical results from the analysis of vibroacoustic signal Bray-Curtis distances between pre-adoption *P. teleius* caterpillars and *M. scabrinodis* ant workers from different populations. a) Generalized linear model (GLM) variable significance. b) Estimated marginal means (EMMs) pairwise comparisons among groups. The first term of each group in the contrast indicates the ant host population; the second term indicates the population of *P. teleius*. NL refers to the Netherlands and PL refers to Poland.

a)

| Variable | d.f. | LR Chisq | p |
|---|---|---|---|
| Worker population | 1 | 0.00 | 0.959 |
| Caterpillar population | 1 | 14.56 | 0.001*** |
| Worker population x caterpillar population | 1 | 2,999.91 | 0.001*** |

*p ≤ 0.05, **p ≤ 0.01, ***p ≤ 0.001

b)

| Contrast | z | p |
|---|---|---|
| NL-NL vs PL-NL | -3.41 | 0.003** |
| NL-NL vs NL-PL | 0.79 | 0.428 |
| NL-NL vs PL-PL | 2.43 | 0.030* |
| PL-NL vs NL-PL | 2.97 | 0.012* |
| PL-NL vs PL-PL | 6.83 | 0.001*** |
| NL-PL vs PL-PL | 2.65 | 0.024* |

*p ≤ 0.05, **p ≤ 0.01, ***p ≤ 0.001

Supplementary material. Sánchez-García, D. et al. Ongoing coevolution between reintroduced *Phengaris teleius* butterflies and their *Myrmica* host ants.

Table S6. Generalized linear mixed model (GLMM) variable significance of different vibroacoustic parameters, i.e., a) peak frequency (Fpeak, Hz), b) third frequency quartile (Q75%, Hz), c) unit duration (Duration, s), d) root-mean-square (RMS, Pa), e) energy of the peak frequency (EnFpeak, Pa2·s), f) percentage of the energy of the peak frequency over the total energy (%EnFpeak, %) of stridulation units of the signals emitted by pre-adopted *P. teleius* caterpillars, *M. scabrinodis* ant queens and *M. scabrinodis* ant workers from the Netherlands and Poland. NL and PL refers to the Netherlands and Poland, respectively, and indicate the population of the ant queens, workers and caterpillars.

a) Peak frequency (Fpeak, Hz)

| Contrast | t | p |
|---|---|---|
| Queen NL vs Worker NL | -4.17 | 0.001*** |
| Queen NL vs P. tel NL | 2.60 | 0.079 |
| Queen NL vs Queen PL | 1.60 | 0.445 |
| Queen NL vs Worker PL | -2.02 | 0.229 |
| Queen NL vs P. tel PL | 2.15 | 0.214 |
| Worker NL vs P. tel NL | 8.09 | 0.001*** |
| Worker NL vs Queen PL | 6.55 | 0.001*** |
| Worker NL vs Worker PL | 3.56 | 0.004** |
| Worker NL vs P. tel PL | 8.08 | 0.001*** |
| P. tel NL vs Queen PL | -1.44 | 0.466 |
| P. tel NL vs Worker PL | -5.74 | 0.001*** |
| P. tel NL vs P. tel PL | -0.66 | 0.759 |
| Queen PL vs Worker PL | -4.04 | 0.001** |

a) Peak frequency (Fpeak, Hz)

| Contrast | t | p |
|---|---|---|
| Queen PL vs P. tel PL | 0.89 | 0.759 |
| Worker PL vs P. tel PL | 5.50 | 0.001*** |

*p ≤ 0.05, **p ≤ 0.01, ***p ≤ 0.001

b) Third frequency quartile (Q75%, Hz)

| Contrast | t | p |
|---|---|---|
| Queen NL vs Worker NL | -6.19 | 0.001*** |
| Queen NL vs P. tel NL | 2.58 | 0.058 |
| Queen NL vs Queen PL | 2.86 | 0.028* |
| Queen NL vs Worker PL | -0.92 | 0.718 |
| Queen NL vs P. tel PL | 4.46 | 0.001*** |
| Worker NL vs P. tel NL | 8.60 | 0.001*** |
| Worker NL vs Queen PL | 10.75 | 0.001*** |
| Worker NL vs Worker PL | 8.22 | 0.001*** |
| Worker NL vs P. tel PL | 11.25 | 0.001*** |
| P. tel NL vs Queen PL | -0.42 | 0.718 |
| P. tel NL vs Worker PL | -3.94 | 0.002** |
| P. tel NL vs P. tel PL | 2.13 | 0.102 |
| Queen PL vs Worker PL | -4.54 | 0.001*** |
| Queen PL vs P. tel PL | 2.40 | 0.081 |
| Worker PL vs P. tel PL | 6.35 | 0.001*** |

*p ≤ 0.05, **p ≤ 0.01, ***p ≤ 0.001

153

c) Unit duration (Duration, s)

| Contrast | t | p |
|---|---|---|
| Queen NL vs Worker NL | 1.75 | 0.674 |
| Queen NL vs P. tel NL | 0.80 | 1.000 |
| Queen NL vs Queen PL | -1.62 | 0.674 |
| Queen NL vs Worker PL | -2.43 | 0.198 |
| Queen NL vs P. tel PL | -0.63 | 1.000 |
| Worker NL vs P. tel NL | -0.94 | 1.000 |
| Worker NL vs Queen PL | -3.70 | 0.009** |
| Worker NL vs Worker PL | -7.38 | 0.001*** |
| Worker NL vs P. tel PL | -3.20 | 0.024* |
| P. tel NL vs Queen PL | -2.27 | 0.253 |
| P. tel NL vs Worker PL | -4.02 | 0.001** |
| P. tel NL vs P. tel PL | -1.70 | 0.674 |
| Queen PL vs Worker PL | -1.06 | 1.000 |
| Queen PL vs P. tel PL | 0.79 | 1.000 |
| Worker PL vs P. tel PL | 2.30 | 0.243 |

*p ≤ 0.05, **p ≤ 0.01, ***p ≤ 0.001

d) Root-mean-square (RMS, Pa)

| Contrast | t | p |
|---|---|---|
| Queen NL vs Worker NL | 3.01 | 0.012* |
| Queen NL vs P. tel NL | -3.12 | 0.011* |
| Queen NL vs Queen PL | 5.71 | 0.001*** |
| Queen NL vs Worker PL | 10.73 | 0.001*** |
| Queen NL vs P. tel PL | 2.83 | 0.016* |
| Worker NL vs P. tel NL | -6.15 | 0.001*** |
| Worker NL vs Queen PL | 3.54 | 0.003** |
| Worker NL vs Worker PL | 12.05 | 0.001*** |
| Worker NL vs P. tel PL | 0.84 | 0.405 |
| P. tel NL vs Queen PL | 7.85 | 0.001*** |
| P. tel NL vs Worker PL | 12.53 | 0.001*** |
| P. tel NL vs P. tel PL | 7.39 | 0.001*** |
| Queen PL vs Worker PL | 5.57 | 0.001*** |
| Queen PL vs P. tel PL | -1.53 | 0.259 |
| Worker PL vs P. tel PL | -5.68 | 0.001*** |

*$p \leq 0.05$, **$p \leq 0.01$, ***$p \leq 0.001$

e) Energy of the peak frequency (EnFpeak, Pa2·s)

| Contrast | t | p |
|---|---|---|
| Queen NL vs Worker NL | 7.36 | 0.001*** |
| Queen NL vs P. tel NL | -2.23 | 0.082 |
| Queen NL vs Queen PL | 5.98 | 0.001*** |
| Queen NL vs Worker PL | 14.21 | 0.001*** |
| Queen NL vs P. tel PL | 2.96 | 0.019* |
| Worker NL vs P. tel NL | -8.76 | 0.001*** |
| Worker NL vs Queen PL | -1.25 | 0.213 |
| Worker NL vs Worker PL | 10.77 | 0.001*** |
| Worker NL vs P. tel PL | -2.76 | 0.031* |
| P. tel NL vs Queen PL | 7.16 | 0.001*** |
| P. tel NL vs Worker PL | 14.54 | 0.001*** |
| P. tel NL vs P. tel PL | 6.40 | 0.001*** |
| Queen PL vs Worker PL | 9.34 | 0.001*** |
| Queen PL vs P. tel PL | -1.67 | 0.197 |
| Worker PL vs P. tel PL | -8.71 | 0.001*** |

*p ≤ 0.05, **p ≤ 0.01, ***p ≤ 0.001

f) Percentage of the energy of the peak frequency over the total energy (%EnFpeak, %)

| Contrast | t | p |
|---|---|---|
| Queen NL vs Worker NL | 9.63 | 0.001*** |
| Queen NL vs P. tel NL | -1.59 | 0.232 |
| Queen NL vs Queen PL | 3.93 | 0.001*** |
| Queen NL vs Worker PL | 11.31 | 0.001*** |
| Queen NL vs P. tel PL | 0.46 | 0.647 |
| Worker NL vs P. tel NL | -10.40 | 0.001*** |
| Worker NL vs Queen PL | -6.47 | 0.001*** |
| Worker NL vs Worker PL | 2.76 | 0.031* |
| Worker NL vs P. tel PL | -8.33 | 0.001*** |
| P. tel NL vs Queen PL | 4.96 | 0.001*** |
| P. tel NL vs Worker PL | 11.96 | 0.001*** |
| P. tel NL vs P. tel PL | 2.49 | 0.040* |
| Queen PL vs Worker PL | 8.46 | 0.001*** |
| Queen PL vs P. tel PL | -2.87 | 0.031* |
| Worker PL vs P. tel PL | -9.97 | 0.001*** |

*p ≤ 0.05, **p ≤ 0.01, ***p ≤ 0.001

Supplementary material. Sánchez-García, D. et al. Ongoing coevolution between reintroduced *Phengaris teleius* butterflies and their *Myrmica* host ants.

Table S7. EMMs (estimated marginal means) test results of the pairwise comparisons of different vibroacoustic parameters, i.e., a) peak frequency (Fpeak, Hz), b) third frequency quartile (Q75%, Hz), c) unit duration (Duration, s), d) root-mean-square (RMS, Pa), e) energy of the peak frequency (EnFpeak, Pa2·s), f) percentage of the energy of the peak frequency over the total energy (%EnFpeak, %) of stridulation units of the signals emitted by pre-adopted *P. teleius* caterpillars, *M. scabrinodis* ant queens and *M. scabrinodis* ant workers from the Netherlands and Poland. NL and PL refers to the Netherlands and Poland, respectively, and indicate the population of the ant queens, workers and caterpillars.

a) Peak frequency (Fpeak, Hz)

| Contrast | t | p |
|---|---|---|
| Queen NL vs Worker NL | -4.17 | 0.001*** |
| Queen NL vs P. tel NL | 2.60 | 0.079 |
| Queen NL vs Queen PL | 1.60 | 0.445 |
| Queen NL vs Worker PL | -2.02 | 0.229 |
| Queen NL vs P. tel PL | 2.15 | 0.214 |
| Worker NL vs P. tel NL | 8.09 | 0.001*** |
| Worker NL vs Queen PL | 6.55 | 0.001*** |
| Worker NL vs Worker PL | 3.56 | 0.004** |
| Worker NL vs P. tel PL | 8.08 | 0.001*** |
| P. tel NL vs Queen PL | -1.44 | 0.466 |
| P. tel NL vs Worker PL | -5.74 | 0.001*** |

a) Peak frequency (Fpeak, Hz)

| Contrast | t | p |
|---|---|---|
| P. tel NL vs P. tel PL | -0.66 | 0.759 |
| Queen PL vs Worker PL | -4.04 | 0.001** |
| Queen PL vs P. tel PL | 0.89 | 0.759 |
| Worker PL vs P. tel PL | 5.50 | 0.001*** |

*$p \leq 0.05$, **$p \leq 0.01$, ***$p \leq 0.001$

b) Third frequency quartile (Q75%, Hz)

| Contrast | t | p |
|---|---|---|
| Queen NL vs Worker NL | -6.19 | 0.001*** |
| Queen NL vs P. tel NL | 2.58 | 0.058 |
| Queen NL vs Queen PL | 2.86 | 0.028* |
| Queen NL vs Worker PL | -0.92 | 0.718 |
| Queen NL vs P. tel PL | 4.46 | 0.001*** |
| Worker NL vs P. tel NL | 8.60 | 0.001*** |
| Worker NL vs Queen PL | 10.75 | 0.001*** |
| Worker NL vs Worker PL | 8.22 | 0.001*** |
| Worker NL vs P. tel PL | 11.25 | 0.001*** |
| P. tel NL vs Queen PL | -0.42 | 0.718 |
| P. tel NL vs Worker PL | -3.94 | 0.002** |
| P. tel NL vs P. tel PL | 2.13 | 0.102 |
| Queen PL vs Worker PL | -4.54 | 0.001*** |
| Queen PL vs P. tel PL | 2.40 | 0.081 |
| Worker PL vs P. tel PL | 6.35 | 0.001*** |

*p ≤ 0.05, **p ≤ 0.01, ***p ≤ 0.001

c) Unit duration (Duration, s)

| Contrast | t | p |
|---|---|---|
| Queen NL vs Worker NL | 1.75 | 0.674 |
| Queen NL vs P. tel NL | 0.80 | 1.000 |
| Queen NL vs Queen PL | -1.62 | 0.674 |
| Queen NL vs Worker PL | -2.43 | 0.198 |
| Queen NL vs P. tel PL | -0.63 | 1.000 |
| Worker NL vs P. tel NL | -0.94 | 1.000 |
| Worker NL vs Queen PL | -3.70 | 0.009** |
| Worker NL vs Worker PL | -7.38 | 0.001*** |
| Worker NL vs P. tel PL | -3.20 | 0.024* |
| P. tel NL vs Queen PL | -2.27 | 0.253 |
| P. tel NL vs Worker PL | -4.02 | 0.001** |
| P. tel NL vs P. tel PL | -1.70 | 0.674 |
| Queen PL vs Worker PL | -1.06 | 1.000 |
| Queen PL vs P. tel PL | 0.79 | 1.000 |
| Worker PL vs P. tel PL | 2.30 | 0.243 |

*p ≤ 0.05, **p ≤ 0.01, ***p ≤ 0.001

d) Root-mean-square (RMS, Pa)

| Contrast | t | p |
|---|---|---|
| Queen NL vs Worker NL | 3.01 | 0.012* |
| Queen NL vs P. tel NL | -3.12 | 0.011* |
| Queen NL vs Queen PL | 5.71 | 0.001*** |
| Queen NL vs Worker PL | 10.73 | 0.001*** |
| Queen NL vs P. tel PL | 2.83 | 0.016* |
| Worker NL vs P. tel NL | -6.15 | 0.001*** |
| Worker NL vs Queen PL | 3.54 | 0.003** |
| Worker NL vs Worker PL | 12.05 | 0.001*** |
| Worker NL vs P. tel PL | 0.84 | 0.405 |
| P. tel NL vs Queen PL | 7.85 | 0.001*** |
| P. tel NL vs Worker PL | 12.53 | 0.001*** |
| P. tel NL vs P. tel PL | 7.39 | 0.001*** |
| Queen PL vs Worker PL | 5.57 | 0.001*** |
| Queen PL vs P. tel PL | -1.53 | 0.259 |
| Worker PL vs P. tel PL | -5.68 | 0.001*** |

*p ≤ 0.05, **p ≤ 0.01, ***p ≤ 0.001

e) Energy of the peak frequency (EnFpeak, Pa2·s)

| Contrast | t | p |
|---|---|---|
| Queen NL vs Worker NL | 7.36 | 0.001*** |
| Queen NL vs P. tel NL | -2.23 | 0.082 |
| Queen NL vs Queen PL | 5.98 | 0.001*** |
| Queen NL vs Worker PL | 14.21 | 0.001*** |
| Queen NL vs P. tel PL | 2.96 | 0.019* |
| Worker NL vs P. tel NL | -8.76 | 0.001*** |
| Worker NL vs Queen PL | -1.25 | 0.213 |
| Worker NL vs Worker PL | 10.77 | 0.001*** |
| Worker NL vs P. tel PL | -2.76 | 0.031* |
| P. tel NL vs Queen PL | 7.16 | 0.001*** |
| P. tel NL vs Worker PL | 14.54 | 0.001*** |
| P. tel NL vs P. tel PL | 6.40 | 0.001*** |
| Queen PL vs Worker PL | 9.34 | 0.001*** |
| Queen PL vs P. tel PL | -1.67 | 0.197 |
| Worker PL vs P. tel PL | -8.71 | 0.001*** |

*p ≤ 0.05, **p ≤ 0.01, ***p ≤ 0.001

173

f) Percentage of the energy of the peak frequency over the total energy (%EnFpeak, %)

| Contrast | t | p |
|---|---|---|
| Queen NL vs Worker NL | 9.63 | 0.001*** |
| Queen NL vs P. tel NL | -1.59 | 0.232 |
| Queen NL vs Queen PL | 3.93 | 0.001*** |
| Queen NL vs Worker PL | 11.31 | 0.001*** |
| Queen NL vs P. tel PL | 0.46 | 0.647 |
| Worker NL vs P. tel NL | -10.40 | 0.001*** |
| Worker NL vs Queen PL | -6.47 | 0.001*** |
| Worker NL vs Worker PL | 2.76 | 0.031* |
| Worker NL vs P. tel PL | -8.33 | 0.001*** |
| P. tel NL vs Queen PL | 4.96 | 0.001*** |
| P. tel NL vs Worker PL | 11.96 | 0.001*** |
| P. tel NL vs P. tel PL | 2.49 | 0.040* |
| Queen PL vs Worker PL | 8.46 | 0.001*** |
| Queen PL vs P. tel PL | -2.87 | 0.031* |
| Worker PL vs P. tel PL | -9.97 | 0.001*** |

*p ≤ 0.05, **p ≤ 0.01, ***p ≤ 0.001

Supplementary material. Sánchez-García, D. et al. Ongoing coevolution between reintroduced *Phengaris teleius* butterflies and their *Myrmica* host ants.

Table S8. Statistical results from the analysis of ant responses to different vibroacoustic signals during playback experiments. a) Generalized linear model (GLM) variable significance of ant worker data from Poland. b) Estimated marginal means (EMMs) pairwise comparisons among responses of ants from Poland to different vibroacoustic signals. c) Generalized linear model (GLM) variable significance of ant worker data from the Netherlands. d) Estimated marginal means (EMMs) pairwise comparisons among responses of ants from the Netherlands to different vibroacoustic signals. Queen refers to the ant queen signal, worker refers to the ant worker signal and P. tel refers to the signal produced by *P. teleius* caterpillars. NL and PL refers to the Netherlands and Poland, respectively, and indicate the population of origin.

a) GLM variable significance of ant workers from Poland

| Variable | d.f. | Chisq | p |
|---|---|---|---|
| Vibroacoustic signal | 4 | 404.66 | 0.001*** |

*p ≤ 0.05, **p ≤ 0.01, ***p ≤ 0.001

b) EMMs pairwise comparison of ant workers from Poland

| Contrast | z | p |
|---|---|---|
| Queen vs P. tel PL | 13.81 | 0.001*** |
| Queen vs P. tel NL | 10.09 | 0.001*** |
| Queen vs White noise | 15.18 | 0.001*** |
| Queen vs Worker | 8.38 | 0.001*** |

b) EMMs pairwise comparison of ant workers from Poland

| Contrast | z | p |
|---|---|---|
| P. tel PL vs P. tel NL | -5.08 | 0.001*** |
| P. tel PL vs White noise | 6.04 | 0.001*** |
| P. tel PL vs Worker | -6.86 | 0.001*** |
| P. tel NL vs White noise | 9.73 | 0.001*** |
| P. tel NL vs Worker | -1.94 | 0.520 |
| White noise vs Worker | -10.91 | 0.001*** |

*p ≤ 0.05, **p ≤ 0.01, ***p ≤ 0.001

188

c) GLM variable significance of ant workers from the Netherlands

| Variable | d.f. | Chisq | p |
|---|---|---|---|
| Vibroacoustic signal | 4 | 49.52 | 0.001*** |

*p ≤ 0.05, **p ≤ 0.01, ***p ≤ 0.001

d) EMMs pairwise comparison of ant workers from the Netherlands

| Contrast | z | p |
|---|---|---|
| Queen vs P. tel PL | -1.35 | 1.000 |
| Queen vs P. tel NL | 2.98 | 0.028* |
| Queen vs White noise | 4.81 | 0.001*** |
| Queen vs Worker | -0.38 | 1.000 |
| P. tel PL vs P. tel NL | 4.32 | 0.001*** |
| P. tel PL vs White noise | 6.05 | 0.001*** |
| P. tel PL vs Worker | 0.96 | 1.000 |
| P. tel NL vs White noise | 2.01 | 0.448 |
| P. tel NL vs Worker | -3.33 | 0.009** |
| White noise vs Worker | -5.13 | 0.001*** |

*p ≤ 0.05, **p ≤ 0.01, ***p ≤ 0.001

Supplementary material. Sánchez-García, D. et al. Ongoing coevolution between reintroduced *Phengaris teleius* butterflies and their *Myrmica* host ants.

Table S9. Statistical results from the analysis of the number of antennations produced by *M. scabrinodis* ant workers in the presence of *P. teleius* caterpillars from the Netherlands and Poland during the behavioral experiment. a) Generalized linear model (GLM) variable significance. b) Estimated marginal means (EMMs) pairwise comparisons. The first term of each group in the contrast indicates the ant host population; the second term indicates the population of *P. teleius*. NL refers to the Netherlands and PL refers to Poland.

a) GLM variable significance

| Variable | d.f. | Chisq | p |
|---|---|---|---|
| Ant population | 1 | 2.97 | 0.085 |
| Caterpillar population | 1 | 4.74 | 0.030* |
| Ant population x Caterpillar population | 1 | 0.69 | 0.405 |

*p ≤ 0.05, **p ≤ 0.01, ***p ≤ 0.001

b) EMMs pairwise comparisons

| Contrast | Z | p |
|---|---|---|
| NL-NL vs PL-NL | -0.50 | 1.000 |
| NL-NL vs NL-PL | -0.70 | 1.000 |
| NL-NL vs PL-PL | -2.23 | 0.153 |
| PL-NL vs NL-PL | -0.29 | 1.000 |
| PL-NL vs PL-PL | -2.29 | 0.132 |
| NL-PL vs PL-PL | -1.87 | 0.371 |

b) EMMs pairwise comparisons

| Contrast | Z | p |
|---|---|---|

*p ≤ 0.05, **p ≤ 0.01, ***p ≤ 0.001

Supplementary material. Sánchez-García, D. et al. Ongoing coevolution between reintroduced *Phengaris teleius* butterflies and their *Myrmica* host ants.

Table S10. Statistical results from the analysis of the number of positive behaviors produced by *M. scabrinodis* ant workers in the presence of *P. teleius* caterpillars from the Netherlands and Poland during the behavioral experiment. a) Generalized linear model (GLM) variable significance. b) Estimated marginal means (EMMs) pairwise comparisons. The first term of each group in the contrast indicates the ant host population; the second term indicates the population of *P. teleius*. NL refers to the Netherlands and PL refers to Poland.

a) GLM variable significance

| Variable | d.f. | Chisq | p |
|---|---|---|---|
| Ant population | 1 | 0.38 | 0.540 |
| Caterpillar population | 1 | 0.97 | 0.325 |
| Ant population x Caterpillar population | 1 | 8.18 | 0.004** |

*p ≤ 0.05, **p ≤ 0.01, ***p ≤ 0.001

b) EMMs pairwise comparisons

| Contrast | Z | p |
|---|---|---|
| NL-NL vs PL-NL | 2.56 | 0.063 |
| NL-NL vs NL-PL | 1.30 | 1.000 |
| NL-NL vs PL-PL | -0.05 | 1.000 |
| PL-NL vs NL-PL | -1.27 | 1.000 |
| PL-NL vs PL-PL | -2.74 | 0.037* |
| NL-PL vs PL-PL | -1.43 | 0.918 |

b) EMMs pairwise comparisons

| Contrast | Z | p |
|---|---|---|

*p ≤ 0.05, **p ≤ 0.01, ***p ≤ 0.001

Supplementary material. Sánchez-García, D. et al. Ongoing coevolution between reintroduced *Phengaris teleius* butterflies and their *Myrmica* host ants.

Table S11. Statistical results from the analysis of the number of negative behaviors produced by *M. scabrinodis* ant workers in the presence of *P. teleius* caterpillars from the Netherlands and Poland during the behavioral experiment. a) Generalized linear model (GLM) variable significance. b) Estimated marginal means (EMMs) pairwise comparisons. The first term of each group in the contrast indicates the ant host population; the second term indicates the population of *P. teleius*. NL refers to the Netherlands and PL refers to Poland.

a) GLM variable significance

| Variable | d.f. | Chisq | p |
|---|---|---|---|
| Ant population | 1 | 5.62 | 0.018* |
| Caterpillar population | 1 | 3.03 | 0.082 |
| Ant population x Caterpillar population | 1 | 4.51 | 0.034* |

*p ≤ 0.05, **p ≤ 0.01, ***p ≤ 0.001

b) EMMs pairwise comparisons

| Contrast | Z | p |
|---|---|---|
| NL-NL vs PL-NL | 0.25 | 1.000 |
| NL-NL vs NL-PL | -0.40 | 1.000 |
| NL-NL vs PL-PL | 2.76 | 0.035* |
| PL-NL vs NL-PL | -0.68 | 1.000 |
| PL-NL vs PL-PL | 2.72 | 0.039* |
| NL-PL vs PL-PL | 3.17 | 0.009** |

b) EMMs pairwise comparisons

| Contrast | Z | p |
|---|---|---|

*p ≤ 0.05, **p ≤ 0.01, ***p ≤ 0.001

Supplementary material. Sánchez-García, D. et al. Ongoing coevolution between reintroduced *Phengaris teleius* butterflies and their *Myrmica* host ants.

Table S12. Statistical results from the analysis of the number of drops produced by *P. teleius* caterpillars from different metapopulations and exposed to different host ants. a) Generalized linear model (GLM) variable significance. b) Estimated marginal means (EMMs) pairwise comparison. The first term of each group in the contrast indicates the ant host population; the second term indicates the population of *P. teleius*. NL refers to the Netherlands and PL refers to Poland.

a) GLM variable significance

| Variable | d.f. | Chisq | p |
|---|---|---|---|
| Ant population | 1 | 0.04 | 0.848 |
| Caterpillar population | 1 | 0.00 | 0.944 |
| Ant population x Caterpillar population | 1 | 0.77 | 0.379 |

*p ≤ 0.05, **p ≤ 0.01, ***p ≤ 0.001

b) EMMs pairwise comparisons

| Contrast | Z | p |
|---|---|---|
| NL-NL vs PL-NL | 0.48 | 1.000 |
| NL-NL vs NL-PL | 0.63 | 1.000 |
| NL-NL vs PL-PL | -0.09 | 1.000 |
| PL-NL vs NL-PL | 0.19 | 1.000 |
| PL-NL vs PL-PL | -0.62 | 1.000 |
| NL-PL vs PL-PL | -0.76 | 1.000 |

*p ≤ 0.05, **p ≤ 0.01, ***p ≤ 0.001

Supplementary material. Sánchez-García, D. et al. Ongoing coevolution between reintroduced *Phengaris teleius* butterflies and their *Myrmica* host ants.

Table S13. Statistical results from the analysis of the adoption proportion of *P. teleius* caterpillars in the presence of *M. scabrinodis* ants from the Netherlands and Poland. a) Generalized linear model (GLM) variable significance. b) Estimated marginal means (EMMs) pairwise comparison. The first term of each group in the contrast indicates the ant host population; the second term indicates the population of *P. teleius*. NL refers to the Netherlands and PL refers to Poland.

a) GLM variable significance

| Variable | d.f. | Chisq | p |
|---|---|---|---|
| Ant population | 1 | 1.68 | 0.195 |
| Caterpillar population | 1 | 4.13 | 0.042* |

*p ≤ 0.05, **p ≤ 0.01, ***p ≤ 0.001

b) EMMs pairwise comparisons

| Contrast | Z | p |
|---|---|---|
| NL-NL vs PL-NL | -1.30 | 1.000 |
| NL-NL vs NL-PL | -2.03 | 0.253 |
| NL-NL vs PL-PL | -2.30 | 0.128 |
| PL-NL vs NL-PL | -0.89 | 1.000 |
| PL-NL vs PL-PL | -2.03 | 0.253 |
| NL-PL vs PL-PL | -1.30 | 1.000 |

*p ≤ 0.05, **p ≤ 0.01, ***p ≤ 0.001

Supplementary material. Sánchez-García, D. et al. Ongoing coevolution between reintroduced *Phengaris teleius* butterflies and their *Myrmica* host ants.

Table S14. Statistical results from the analysis of the survival probability of *P. teleius* caterpillars in the presence of *M. scabrinodis* ants from the Netherlands and Poland. a) Generalized linear model (GLM) variable significance. b) Estimated marginal means (EMMs) pairwise comparison.The first term of each group in the contrast indicates the ant host population; the second term indicates the population of *P. teleius*. NL refers to the Netherlands and PL refers to Poland.

a) GLM variable significance

| Variable | d.f. | LR Chisq | p |
|---|---|---|---|
| Ant population | 1 | 4.04 | 0.044* |
| Caterpillar population | 1 | 3.50 | 0.061 |

*p ≤ 0.05, **p ≤ 0.01, ***p ≤ 0.001

b) EMMs pairwise comparisons

| Contrast | Z | p |
|---|---|---|
| NL-NL vs PL-NL | 2.05 | 0.240 |
| NL-NL vs NL-PL | 1.87 | 0.366 |
| NL-NL vs PL-PL | 2.80 | 0.030* |
| PL-NL vs NL-PL | -0.22 | 1.000 |
| PL-NL vs PL-PL | 1.87 | 0.366 |
| NL-PL vs PL-PL | 2.05 | 0.240 |

*p ≤ 0.05, **p ≤ 0.01, ***p ≤ 0.001

Supplementary material. Sánchez-García, D. et al. Ongoing coevolution between reintroduced *Phengaris teleius* butterflies and their *Myrmica* host ants.

Table S15. Cox proportional-hazard ratios. On the left, the predictor variables (ant population and caterpillar population) and levels (NL: the Netherlands, PL: Poland). N refers to the sample size of each level. In the middle, hazard ratios. The dotted vertical line represents the reference level. NL was taken as a reference level for the two variables. On the right, the value and interval of confidence of the hazard ratio of each variable. p indicates the significance of each variable.

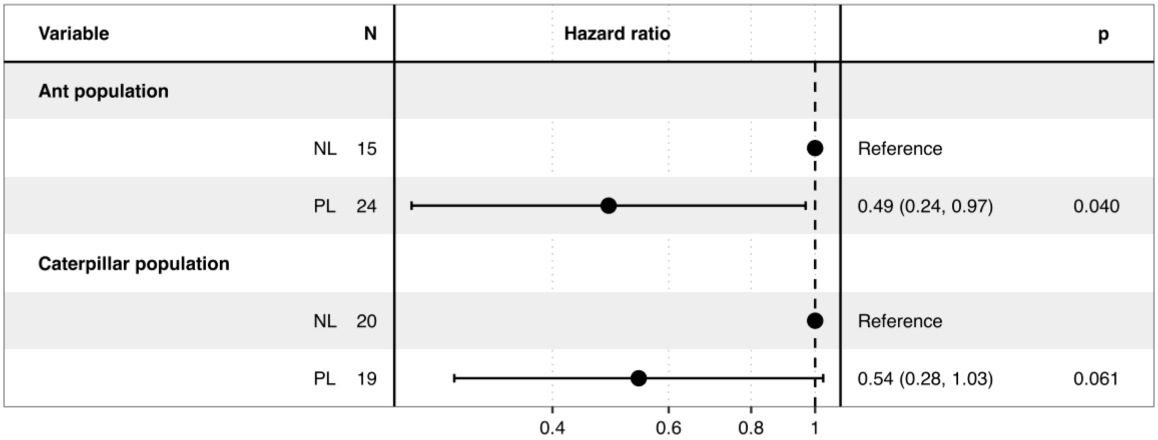
