## Supplementary Methods S1 for "Ongoing coevolution between reintroduced *Phengaris teleius* butterflies and their *Myrmica* host ants"

### Sample collection, transport and maintenance details

*M. scabrinodis* ant colonies used in the cross-population experiment were collected in the field in both populations and transported to Warsaw before the *P. teleius* caterpillars collection. From each ant colony, two sub-colonies consisting of 100 workers (50 foragers and 50 intra-nidal workers) and 30 ant larvae were established in plastic boxes. Sub-colonies of *M. scabrinodis* ants were kept in plastic boxes ($23 \times 15 \times 6$ cm) in which walls were covered with paraffin to prevent workers from escaping. A wet sponge covered by a plastic lid with an entrance notch to provide a suitable and dark place for ants was put on the side. Colonies were fed twice a week with a solution of sugar water and pieces of crickets placed on a circular metallic plate (Ø 3 cm). *P. teleius* caterpillars were collected when the ant colonies were already established in the laboratory. *Sanguisorba officinalis* single stems bearing *P. teleius* caterpillars were collected in the Netherlands and shipped to the laboratory in Warsaw within 24 hours of collection. Similarly, plants with caterpillars collected in the Polish population were transported to the laboratory in Warsaw on the day of collection. These plants were gathered into bunches of a few stems, placing the base of the stems in water. Each bunch was kept in a plastic container, in which the walls were covered by fluon to prevent *P. teleius* caterpillars from escaping. The containers were checked every morning and late afternoon to obtain butterfly caterpillars, and only the fourth-instar caterpillars were used in the study.

### CHC GC-MS analysis details

For shipping, samples were evaporated. Prior to chemical analyses, *P. teleius* caterpillar samples were suspended in a final volume of $20\,\mu L$ of pentane (HPLC grade; Sigma Aldrich) with n-heptadecane (n-$C_{17}$: at $5\,ng/\mu L$ in $100\,ng$) as the internal standard. The *M. scabrinodis* ant extracts were suspended in $60\,\mu L$ of pentane, with n-eicosane (n-$C_{20}$: at $5\,ng/\mu L$ in $300\,ng$) as the internal standard. For caterpillar samples, 3 µL were added in manual injection, and for ant worker samples, an aliquot of $2\ \mu L$ of the solution was injected using an Agilent G4513A Automatic Liquid Sampler into an Agilent Technologies 7890A gas chromatograph coupled with an Agilent 5975 C mass spectrometer (Agilent Technologies, Les Ulis, France). The GC column HP5MS Agilent Technologies ($30\ m \times 0.25\ mm \times 0.25\ \mu m$ film) was coated with (5%-Phenyl)-methylpolysiloxane, and helium was used as a carrier gas ($1\ mL/min$). The injection was splitless, and the oven temperature was set at $60\ °C$ for $1\ min$, then it was raised from $60\ °C$ to $220\ °C$ at $20\ °C \cdot min^{-1}$, then to $250\ °C$ at $3\ °C \cdot min^{-1}$ and then to $5\ °C \cdot min^{-1}$ and held for $5\ min$. Mass spectra were recorded with electron impact ionization at $70$ eV.

**CHC raw data processing details**

Peak integration was performed with the RTE integrator applying the next parameter values: Data point sampling = 1, detection filtering = 7 points, start threshold = 0.020, stop threshold = 0.000, baseline reset > 50, if leading or trailing edge < 100, minimum peak area = 0.1 % of largest peak, peak location = centroid, maximum number of peaks = 250 and baseline preference = baseline drop else tangent.

**Vibroacoustic signal processing and variable definitions**

We carefully examined segments containing acoustic recordings and digitally saved them in WAV format with a 16-bit amplitude resolution using Audacity version 3.0.3. We selected three sequences of vibroacoustic signals per individual. In ant stridulations, each sequence consisted of five pairs of alternating units ($a$ and $b$), while in caterpillars, each sequence consisted of five units of the same type. Using Praat version 6.2.14, we measured six temporal, spectral, and intensity vibroacoustic variables for each unit: the peak frequency (Fpeak, Hz), the third quartile of the energy spectrum (Q75%, Hz; representing 75% of the call's energy), the unit duration (Δt, s), the mean intensity of the entire call, quantified by the root-mean-square signal level (RMS, dB), the energy of the peak frequency (EnFpeak, $Pa^2 \cdot s$) and the ratio of the peak frequency energy to the total call energy, expressed as a percentage (%EFpeak).

**Adoption assay procedures and behavioral definitions**

To observe adoption, a caterpillar was put on a circular metallic plate (Ø 3 cm) at the furthest distance from the entrance of the ant nest. The time of caterpillar insertion and the time of the first contact with ants were noted. During the initial 60 minutes observations, all behavioral events displayed by *M. scabrinodis* workers were recorded and categorized as inspection (antennation); positive behaviors (grooming, licking larval secretions, and picking up, where the caterpillar was taken by an ant and transported for some distance); or negative behaviors (mandible gaping, biting, and stinging). If adoption did not occur within the first 60 minutes, the boxes were checked every hour for the following five hours. If adoption did not occur during this time, we checked the colony after 24 hours; if the caterpillar was present inside the ant nest (together with workers and brood), we considered that the adoption happened within 24 hours. The survival of the caterpillar was checked every day, starting 24 hours after the adoption until its death, and every week, 10 ant larvae were added to each ant sub-colony to provide a food source for the *P. teleius* caterpillars.
